# Peroxisome import stress impairs ribosome biogenesis and induces integrative stress response through eIF2α phosphorylation

**DOI:** 10.1101/2020.11.19.390609

**Authors:** Kerui Huang, Jinoh Kim, Pham Vo, Ting Miao, Hua Bai

## Abstract

Peroxisome biogenesis diseases (PBDs) are characterized by global defects in peroxisomal function and can result in severe brain, liver, kidney, and bone malfunctions. PBDs are due to mutations in peroxisome biogenesis factors (PEX genes) that are responsible for peroxisome assembly and function. Increasing evidence suggests that peroxisome import functions decline during aging. However, the transcriptome profiling of peroxisome import defects and how they affect disease development are still lacking. *PEX5* encodes the cytoplasmic receptors for peroxisome-targeting signal types 1. We generate knock-in human HEK293 cells mutant using CRISPR to transiently express *PEX5* cysteine 11 to alanine mutant (PEX5^C11A^), which blocks PEX5 recycling and exerts dominant negative effect on *PEX5* mediated peroxisome import. To identify conserved responses, we perform transcriptomic analysis on *Drosophila* oenocyte-specific Pex1, Pex12 and Pex5 knockdowns and on human cells with impaired peroxisome import (PEX5^C11A^ and PEX5 siRNA respectively). PEX5^C11A^ induction triggers vast transcriptomic changes, including decreased oxidative phosphorylation, increased MAPK signaling and HIPPO signaling. PEX5 siRNA specifically decreases spliceosome activity and increases cholesterol metabolism. Using gene set enrichment analysis (GSEA), we identify protein processing in endoplasmic reticulum pathway, specifically ER-associated protein degradation (ERAD) pathway is induced in all PEX knockdowns in *Drosophila*. Peroxisome dysfunction elevates eIF2α phosphorylation in both *Drosophila* and human cell culture independent of *XBP1* activation, suggesting increased integrative stress response (ISR). Moreover, peroxisome stress decreases ribosome biogenesis genes and impairs ribosome biogenesis in flies and human cells. Specifically, peroxisome stress impairs the 5’-ETS cleavage activity during the ribosome biogenesis and dampens 40S small ribosomal export in both flies and human. Our results suggest that reduced ribosome biogenesis and elevated ISR could be conserved cellular response to peroxisome import stress.

## Introduction

Peroxisomes are single-membrane organelles found in almost all eukaryotic cells. Peroxisomes are highly dynamic and versatile, as their composition and activity varies between organisms, different cell types and under different environmental stresses^1^. Generally peroxisomes are responsible for metabolizing branched-chain fatty acids (BCFA) or very long chain fatty acids (VLCFAs), purine catabolism, synthesis of plasmalogens, ether-lipids and bile acids, regulation of hydrogen peroxide and other reactive oxygen species (ROS) ^1, 2^. In mammals, β-oxidation occurs largely in the mitochondria. Peroxisomes are normally required for catabolism of BCFA and VLCFAs, however it can perform β-oxidation on other fatty acids when mitochondria are compromised ^3^. Peroxisomes have close interaction with many other organelles. Peroxisomes can physically interact with other organelles including the endoplasmic reticulum (ER), mitochondria, lipid droplets, lysosomes and chloroplasts, to perform specific functions ^4^.

Peroxisomes are formed by the action of 14 peroxisome assembly factors (Peroxins). The majority of which are involved in translocation of peroxisomal enzymes into the peroxisome matrix. Others are responsible for transporting peroxisome membrane proteins^5–8^. Unlike mitochondria, the peroxisome is composed of single membrane and it could import fully folded and oligomeric proteins into the matrix ^9^. Peroxisomal matrix proteins contain one of two types of intrinsic peroxisomal targeting signals (PTSs), PTS1 or PTS2, which can direct import into the organelle ^9^. Most matrix proteins are targeted through the PTS1 pathway. PTS1 consists of a tripeptide (-SKL or conserved variant) at the C-terminal and it can be recognized by a soluble receptor, Pex5 ^9^. Pex7 will recognize PTS2 containing proteins near the N terminus. Pex5 and Pex7 bind their respective PTSs in the cytosol, and the receptor-cargo complexes will dock to the peroxisome surface, containing Pex13 and Pex14 docking complexes. Cargo is released from Pex5 and imported into peroxisomes. Then Pex5 is mono-ubiquitylated by Pex4 and Pex12, enables Pex5 recycling back to the cytosol for another round of import. Pex5 can also undergo polyubiquitylation (by the ubiquitin-conjugating and ligase enzymes, Ubc4 and Pex2 respectively), which directs Pex5 to the proteasome for degradation (see excellent review in ^10^). Mutations in these peroxin genes and peroxisome resident enzymes can lead to a variety of disorders, known as peroxisome biogenesis disorders (PBD) with involvement of kidney, brain, bone and liver, and death in infants ^11–14^.

Recently there are increasing evidence placing peroxisome, especially peroxisomal import function, as an important regulator for aging. Several studies suggest that peroxisomal import function declines with age ^15–17^. Consistently, our translatomic study shows that the majority of peroxisome genes are downregulated in aged fly oenocytes ^18^. Our previous study identified that peroxisome import activities in fly oenocytes can non-autonomously regulate cardiac arrhythmia during aging ^19^. Several other studies have also implicated that peroxisome is involved in longevity^20–22^. However, it remains elusive how impaired peroxisome import activity contributes to cellular processes such as aging.

Ribosome biogenesis is one of the regulators of aging. The reduced level of ribosomal genes, ribosome biogenesis factors or nutrient sensing pathways (such as TOR signaling), which stimulate ribosome production, can increase the lifespan in multiple organisms, including *C. elegans, D. melanogaster,* yeast, mice and human ^23^. Because the rate of protein translation is proportional to the rate of ribosome biogenesis ^24, 25^, it was suggested that reduced ribosome biogenesis can reduce protein synthesis thus maintaining global proteostasis ^23^. The biogenesis of the 60S and 40S ribosomal subunits follows a complicated pathway in eukaryotic cells, which requires the assembly of four rRNAs and ~80 ribosomal proteins (RPs) ^26^. Ribosome biogenesis occurs in the nucleolus and is initiated by transcription of a large precursor rRNA (pre-rRNA) through RNA polymerase I (Pol I), from which the mature 18S, 5.8S, and 25S rRNAs are generated ^27^. The 5S pre-rRNA is transcribed by RNA polymerase III and incorporated into nascent pre-ribosome in later step ^28^. The pre-rRNA will assemble co-transcriptionally with numerous trans-acting factors and early binding ribosomal proteins for the small subunit, forming a macromolecular complex, named 90S pre-ribosome or small-subunit (SSU) processome ^29, 30^. In the firs 90S assembly intermediate, the pre-rRNA undergoes site-specific base modifications at conserved sites and cleavage reactions. These early cleavages will produce 20S pre-rRNA (precursor to 18S), which becomes part of the early pre-40S particle. The pre-40S particle will follow a simple maturation route to the cytoplasm where mature 40S is produced. Ribosome biogenesis is tightly controlled by nutrient availability and extracellular conditions. Myc and TOR pathways are known regulators of ribosome biogenesis ^31, 32^, however it remains elusive whether there are additional regulators of this complex biological process.

Using human HEK293 and HepG2 cell lines, in combination of *Drosophila* tissue specific RNAi, we sought to characterize conserved cellular responses under peroxisome import stress. Our study discovered distinct transcriptional responses from knocking down different components involved in peroxisome import pathways, suggesting PEXs proteins perform additional activities besides facilitating Pex5. Genes participate in protein process and in endoplasmic-reticulum-associated protein degradation (ERAD) are highly induced across all PEXs knockdowns in flies, suggesting an induced ER stress under peroxisome defects. We also identified repression of oxidative phosphorylation genes and induced inflammation pathways across fly and humans, suggesting close interplay of peroxisome and mitochondria. Finally, for the first time our study discovered a conserved reduction of ribosome biogenesis genes and reduced ribosome biogenesis activity, under peroxisome import stress across fly and human. Together, these findings shed lights on mechanisms of PBD pathology and aging.

## Results

### Transcriptomic analysis for peroxisomal stress response in *Drosophila* oenocytes and human cells

Despite the emerging importance of peroxisome in the regulation of aging and metabolism, little is known how cells mount defensive response to maintain cellular homeostasis. Oenocytes are hepatocyte-like cells that are highly enriched with peroxisomes ^19, 33^. In addition, the oenocyte peroxisome was identified as an important regulator for age-related production of inflammatory factor, unpaired 3 (upd3) ^19^, which could dampen cardiac function non-autonomously. To study cellular responses induced by peroxisomal stress, we performed transcriptomic analysis from *Drosophila* oenocytes and human cell cultures. We target on peroxisome import stress by knocking down key factors (peroxins) involved in peroxisomal import process (Pex5, Pex12 and Pex1) specifically in oenocytes, which were shown to efficiently induce the ROS level and upd3 production ^19^. Pex5 is one of the major peroxisome protein import receptors, which can bind cargo proteins containing peroxisomal targeting signal type 1 (PTS1) and deliver them to peroxisomal matrix through Pex13/Pex14 docking complex. After releasing its cargo in peroxisomal matrix, Pex5 will be poly- or mono-ubiquitinated through complex Pex2/Pex10/Pex12. Monoubiquitylated receptor is then released from the membrane by the ATPases of the ATPases associated with diverse cellular activities (AAA) family, Pex1 and Pex6 ^34^ (Fig. 1A). In the cytosol, the ubiquitin moiety is removed and Pex5 becomes available for another round of import. The polyubiquitylated Pex5 receptor is released from peroxisome following the same way as the monoubiquitylated receptor, except that it is directed for proteasomal degradation ^34^. Knocking down these genes have been shown to greatly impair peroxisome import in oenocytes and S2 cells ^19, 35^. We utilized oenocyte-specific GeneSwitch driver (*PromE*^*GS*^-*Gal4*) ^19^ to transiently knock down peroxin genes in adult stage.

**Figure 1.**
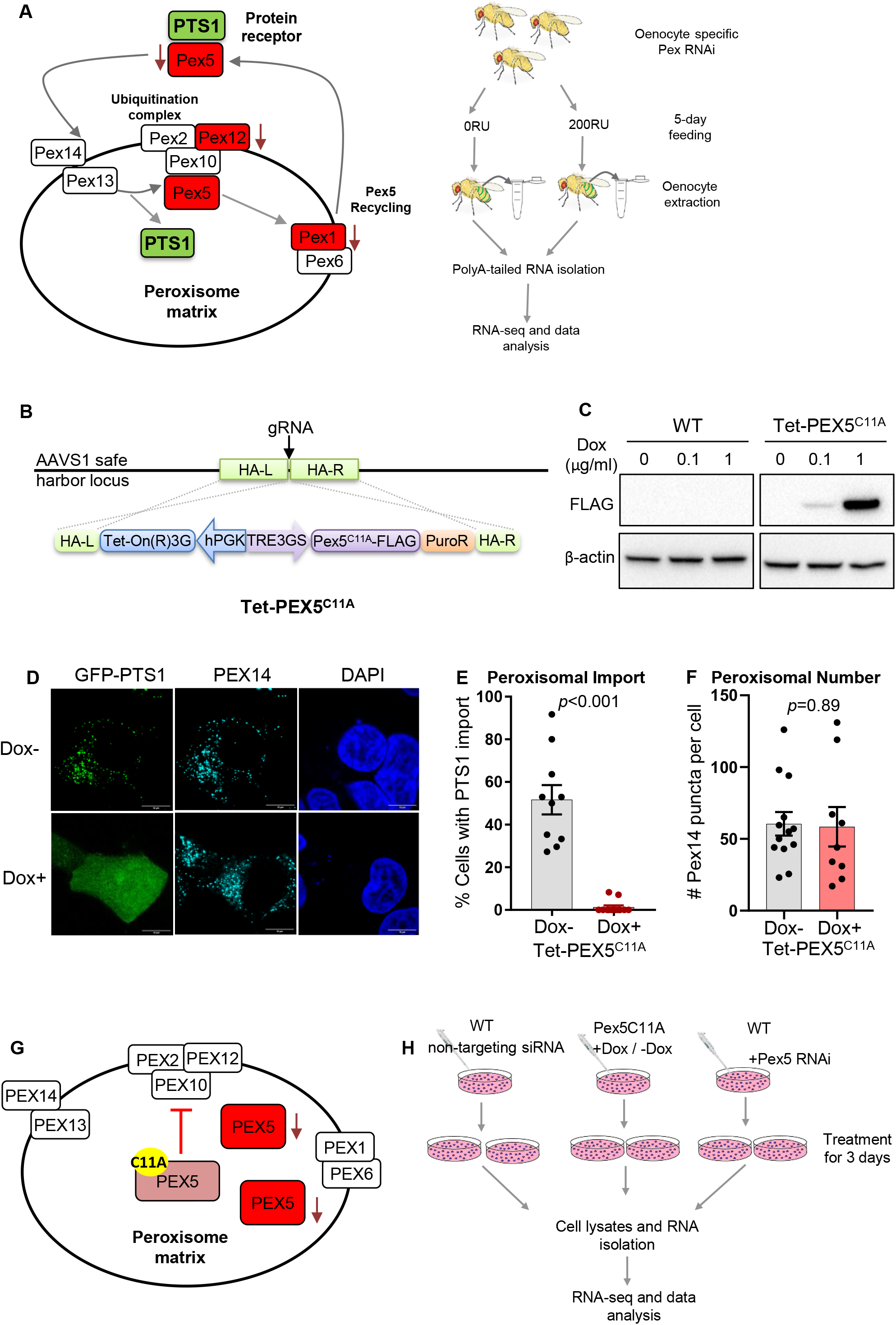
Transcriptomic analysis for peroxisomal stress response in *Drosophila* oenocytes and human cells. **A** Schematic diagram showing the key genes involved in peroxisomal import, as well as oenocyte specific RNA-seq analysis. RNAi targets genes marked in red. **B** Schematic diagram for generation of CRISPR Knock-in HEK293 cells expressing PEX5^C11A^. **C** Western blot shows inducible PEX5^C11A^ construct after treatment of Doxycycline. **D** Immunostaining showing GPF-PTS1 importing activity in wild type cells (Dox-) and PEX5^C11A^ expressing cells (Dox+). Scale bar represents 10μm. **E** Quantification of **D** on % cells with PTS1 import activity in Dox-versus Dox+ cells. **F** Quantification in **D** showing the number of PEX14 positive peroxisomes in Dox-versus Dox+ cells. **G – H** Schematic diagram showing RNA-seq analysis using human cells.

RU486 (mifepristone, or RU) was used to activate *PromE*^*GS*^-*Gal4* (+RU), whereas control genotype is the same, but with no RU feeding (-RU). After 5 days of RU activation, fly oenocytes are dissected and polyA-tailed RNA was isolated for downstream RNA sequencing and data analysis (Fig. 1A). To eliminate the transcriptional changes induced by RU feeding, we included a control group (*PromE-GS-gal4>yw)* for RNA-seq analysis.

To identify conserved pathways involved in peroxisomal stress responses, we have recently developed an inducible peroxisomal stress system in the mammalian cell culture. In this system, we knocked in a Tet-ON 3G tetracycline-inducible expression construct into the AAVS1 safe harbor locus in human embryonic kidney-derived HEK293 cells (Fig. 2B). The expression construct contained a FLAG-tagged full-length human PEX5 with a single amino acid substitution at position 11 (cysteine to alanine, C11A), a conserved ubiquitination site involved in PEX5 recycling ^36^. Stable expression of PEX5^C11A^ mutant exerts dominant-negative effect on wild type PEX5 recycling and efficiently blocks peroxisomal import in mammalian cell culture ^36^. PEX5^C11A^ protein level can be robustly induced by treating cells with doxycycline (Dox) at the concentration of 1 μg/ml, but not in wild type (Fig. 2C). Induction of PEX5^C11A^ mutant was able to block GFP-PTS1 import into peroxisomes (marked by PEX14), without significantly affecting peroxisomal number (Fig. 1D-1F). To conduct RNA-seq analysis, PEX5^C11A^ cells were treated with or without doxycycline for 72 hours prior for total RNA extraction. To directly compare similarity with Pex5 knockdown in *Drosophila* oenocytes, we transfected PEX5-targeting siRNA or non-targeting siRNA into HepG2 cells for 72 hours prior to total RNA isolation. PolyA-tailed RNA was then isolated and RNA-seq libraries were constructed and pooled for the following analysis (Fig. 1G-1H).

**Figure 2.**
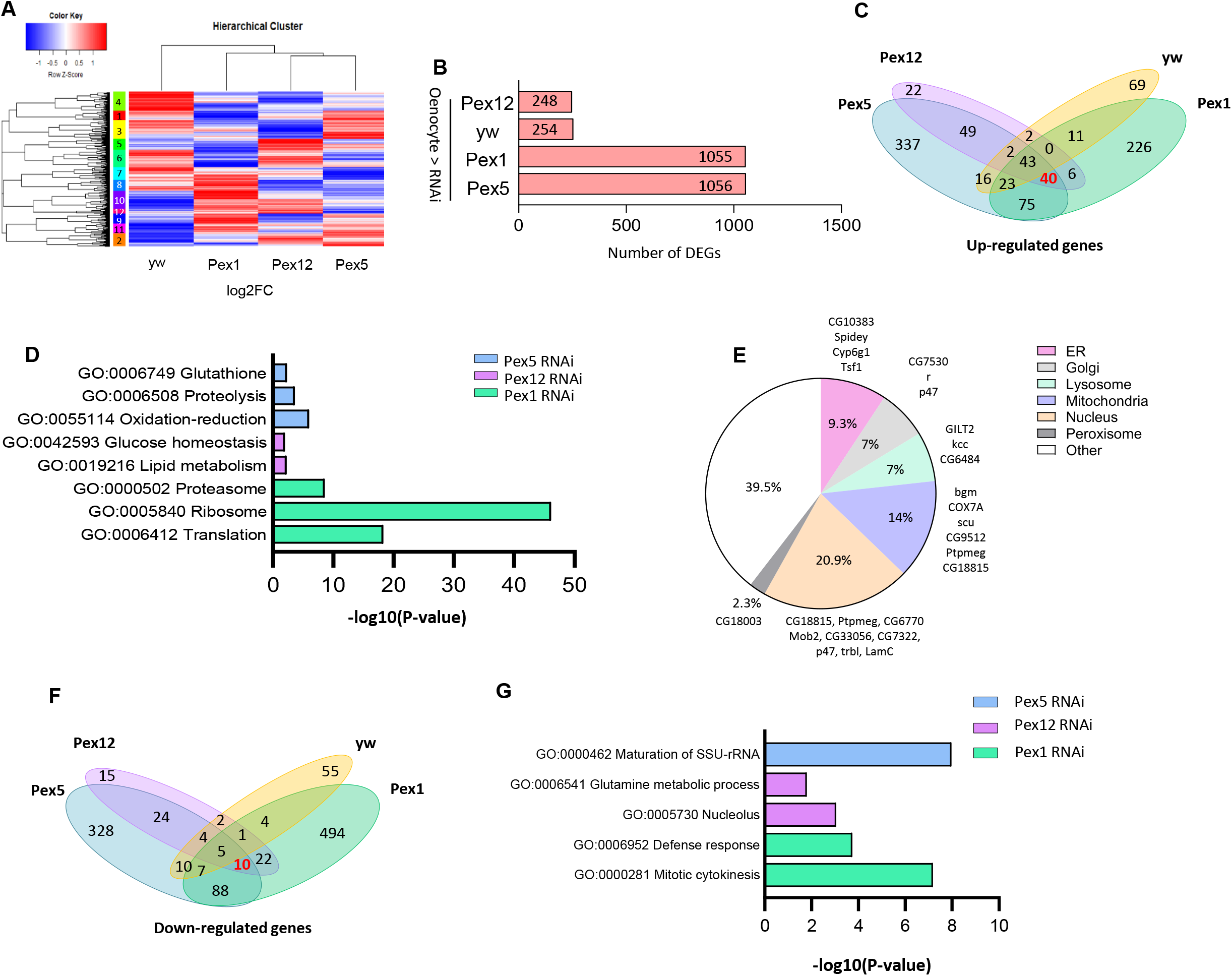
Pex1, Pex12 knock-down show distinct transcriptional pattern from Pex5. **A** Hierarchical clustering heatmap analysis, plotting log2 fold change (+RU / −RU) in yw, Pex1 RNAi, Pex12 RNAi, Pex5 RNAi oenocyte samples. **B** Number of differentially expressed genes (DEGs) across all oenocyte samples. **C** Venn diagram analysis showing genes commonly up regulated by yw, Pex1 RNAi, Pex12 RNAi and Pex5 RNAi (marked by red). **D** Gene Ontology (GO) term analysis on commonly induced genes from **C. E** Organelle localization of proteins produced commonly induced genes. **F** Venn diagram analysis showing genes commonly down regulated by yw, Pex1 RNAi, Pex12 RNAi, and Pex5 RNAi (marked by red). **G** Gene Ontology (GO) term analysis on commonly induced genes from **F**.

#### Distinct transcriptional profiling in Pex1, Pex12 and Pex5 knockdown in oenocytes

To understand how peroxisomal stress induced by different peroxins’ knockdown affect transcriptome, we conducted hierarchical cluster analysis across Pex1^RNAi^, Pex12^RNAi^, Pex5^RNAi^ and yw control samples, using log2 fold change value (RU treatment / no RU treatment) (Fig. 2A). Intriguingly, we found three peroxin knockdowns produced distinct transcriptomic changes with very little similarities, as reflected on the heatmap (Fig. 2A) and PCA analysis (Supplementary Fig. 1A). The transcriptomic profile also varies between PEX5^C11A^ and PEX5-siRNA in human cell culture (Supplementary Fig.1B). The little similarity between each gene knockdowns is possibly due to different methods of inducing peroxisome stress and different cell types used. Cluster 2 in Fig.2A represents genes that are highly induced by both Pex5 and Pex12 and contains GO terms such as response to ER stress, carbohydrate metabolic process, glycogen biosynthetic process and lipid particle (Supplementary Fig 1C). Cluster 4 is enriched with genes specifically induced by RU feeding, which are involved in oogenesis, transcription, histone modification (Supplementary Fig.1D). These results are consistent with previous findings that mifepristone can regulate genes involved in X-chromosome gene expression and oogenesis ^37^. Cluster 9 and 11, which contains genes commonly induced by Pex5 and Pex1, are enriched with GO terms including fatty acid beta-oxidation, mRNA splicing, protein folding, mitochondrial translation, and peroxisome. Induction of peroxisomal genes suggests there is retrograde signaling between peroxisome and nucleus. In cluster 10 and 12 contains genes induced by both Pex1 and Pex12. These clusters contain genes involved in proteasome assembly, translation, mitochondrial electron transport and ribosome Supplementary Fig1E – 1F).

There are 248 differentially expressed genes (DEGs) in Pex12 knock down from oenocytes (│fold change│ ≥ 1.2, FDR ≤ 0.1), 254 DEGs induced by RU feeding. Both Pex5 knockdown and Pex1 knockdown induced great transcriptional changes: Pex1 knockdown produced 1055 DEGs and Pex5 knockdown produced 1056 DEGs (Fig. 2B). Next, we ask what genes are commonly regulated by all Pexs’ knockdowns and what genes are specifically controlled by individual Pexs. We compiled venn diagram using up-regulated DEGs across all Pex-knockdown samples and yw control group. Consistent with the heatmap analysis (Fig.2A), only 40 genes are commonly induced by all Pex knockdowns, but not in yw control group (Fig. 2C), confirming the finding that these Pex knockdowns generate different transcriptional responses. To understand how each Pex genes specifically regulate transcriptions, we conducted GO term analysis on genes specifically induced by individual Pex knock downs. Pex5 knockdown specifically induced 337 DEGs, which are enriched in glutathione, proteolysis, and oxidation-reduction processes. Pex12 knockdown specifically induced genes involved in glucose homeostasis and lipid metabolism. In contrast, Pex1 specifically induced genes involved in protein homeostasis process, such as proteasome, ribosome, and translation (Fig. 2D).

Next, we sought to understand the function of all the commonly induced genes. When we analyzed the function of these genes, we found 9.3% of them function in ER, 7% in Golgi, 7% in lysosome, 14% of them function in mitochondria (Fig. 2E). Majority of the peroxisome stress induced genes produce nucleus-localizing proteins (20.9%), suggesting alterations in nucleus under peroxisome stress. Surprisingly, only 1 peroxisome gene (*CG18003*) is altered by all three Pex knockdowns. Localization predication was based on published datasets on peroxisome and mitochondria ^38, 39^, predicated localization on Flybase and human orthologues predication. From those mitochondrial genes, *COX7A* (cytochrome c oxidase subunit 7A) encodes a subunit for mitochondrial complex IV; CG9512 encodes a glucose-methanol-choline oxidoreductase, its human orthologue CHDH (choline dehydrogenase) localizes to the outer membrane of mitochondria in a potential-dependent manner^40^. CHDH interacts with SQSTM1, a mitophagy adaptor molecule, and CHDH is required for mitophagy ^40^. This suggests mitochondrial abnormalities and possibly mitophagies are induced under peroxisome import stress. Spidey, an oenocyte enriched gene essential for its growth and lipid metabolism, is also induced by peroxisome stress. Its human orthologue (very-long-chain 3-oxoacyl-CoA reductase, HSD17B12) is predicted to localize in the ER. Spidey is reported to be a 3-ketoacyl-CoA reduction for elongation of long chain fatty acids into VLCFAs and is also induced by peroxisome stress^41^ (Fig. 2E). Nucleus genes include *LamC* (*LMNA* in human), *CG6770* (*NUPR1* in human), which regulate nuclear envelope and chromatin organization, are also increased under peroxisome stress. *CG10383 (SERAC1* in human) encodes a hydrolase involved in glycosylphosphatidylinositol metabolism is essential for both mitochondrial function and intracellular cholesterol trafficking. *CG10383* expression increased significantly under all Pex knockdowns (Fig. 2E). *CG7322* is orthologous to *Dicarbonyl/L-xylulose reductase (DCXR)* in human. It encodes a highly conserved enzyme converting L-xylulose into xylitol and it is important in carbonyl detoxification and carbohydrate metabolism ^42^. *DCXR* is thought to play an important role in the glucuronic acid/uronate cycle of glucose metabolism. In this pathway, glucuronic acid is metabolized to form the pentose, L-xylulose. Subsequently, DCXR convers L-xylulose to xylitol which is shuttled into the pentose phosphate pathway. Highly induced expression of *CG7322 (DCXR* orthologue) under peroxisome stress indicates altered pentose phosphate or carbohydrate pathways. We verified the increased expression of *CG10383* and *CG7322* through RT-PCR (Supplementary Fig. 1G-H). Altogether, these data suggest that other organelles’ homeostasis and metabolism are tightly controlled by peroxisomal and these changes can lead collaborative response against disruption of homeostasis.

Comparing to commonly induced genes, Pex knockdowns only commonly repressed the expression of 10 genes (Fig. 2F and Supplementary Fig. 1I-J). Among these 10 genes, 3 of them encode transferases (CG1941, CG2781, CG14615), which can transfer acyl groups other than amino acyl groups. CG2781 (ELOVL fatty acid elongase) is predicated to mediate elongation of fatty acids into very-long-chain fatty acids. Eaat1 (excitatory amino acid transporter 1), which encodes a transmembrane protein with glutamate-sodium symporter activity ^43^, is also commonly repressed by all Pex knockdowns. Its function in oenocytes remain unknown. But glutamate transport could indirectly regulate the synthesis of antioxidant glutathione ^44^, Eaat1 might also regulate glutathione synthesis, thus regulating the redox level in oenocytes. We further analyzed the genes specifically repressed by individual Pex knockdowns. Pex5 knockdown specifically repressed genes involved in maturation of small subunit ribosomal ribonucleic acid (SSU-rRNA). Pex12 knockdown specifically induced glutamine metabolic process and nucleolus genes. Pex1 knockdown repressed genes involved in mitotic cytokinesis, and defense response. Why Pex1 regulates genes in mitotic cytokinesis? It is possibly because they perform other functions in oenocytes, such as cytoskeleton organization. Taken together, our Pex specific transcriptomic analysis revealed common distinct changes elicited by peroxisome stress in adult oenocytes.

### Peroxisomal stress induced endoplasmic reticulum genes and the integrated stress response (ISR)

To further characterize peroxisome-stress induced signaling pathways, we performed Gene Set Enrichment Analysis (GSEA) on three Pex knockdown samples in *Drosophila*, using a collection of pre-defined gene sets retrieved from Kyoto Encyclopedia of Genes and Genomes (KEGG) database on *Drosophila Melanogaster*. Interestingly, Pex12 knockdown affected many pathways involved in RNA metabolism and processing, including RNA polymerase, RNA transport, RNA degradation and spliceosome (Table 1).

**Table 1.** GSEA pathway analysis under Pex5, Pex12 and Pex1 RNAi. NES: normalized enrichment score. “Up or down regulated” indicates whether genes within that pathway are induced or reduced under RNAi treatment.

| NAME | NES | FDR | Type of RNAi | Up or down regulated |
| --- | --- | --- | --- | --- |
| Ribosome biogenesis in eukaryotes | 2.44 | <0.0001 | Pex5 | Down |
| Ribosome | 1.79 | 0.021 | Pex5 | Down |
| DNA replication | 1.68 | 0.06 | Pex5 | Down |
| Protein processing in endoplasmic reticulum | -1.77 | 0.05 | Pex5 | Up |
| Ribosome biogenesis in eukaryotes | 2.89 | <0.0001 | Pex12 | Down |
| DNA replication | 2.11 | <0.0001 | Pex12 | Down |
| RNA polymerase | 1.91 | 0.001 | Pex12 | Down |
| RNA transport | 1.85 | 0.002 | Pex12 | Down |
| Spliceosome | 1.78 | 0.005 | Pex12 | Down |
| RNA degradation | 1.67 | 0.002 | Pex12 | Down |
| Proteasome | -1.90 | 0.01 | Pex12 | Up |
| Protein processing in endoplasmic reticulum | -1.46 | 0.188 | Pex12 | Up |
| TOLL and IMD signaling pathway | 1.74 | 0.041 | Pex1 | Down |
| DNA replication | 1.70 | 0.034 | Pex1 | Down |
| Ribosome | -2.41 | <0.0001 | Pex1 | Up |
| Proteasome | -1.84 | 0.012 | Pex1 | Up |
| Protein processing in endoplasmic reticulum | -1.54 | 0.190 | Pex1 | Up |
| Citrate cycle (TCA-cycle) | -1.43 | 0.295 | Pex1 | Up |

GSEA results showed all three Pex knockdown induced genes involved in DNA replication, which be because of disrupted reactive oxygen species (ROS) level, because ROS are well-known mediators of DNA and mitochondrial DNA damage^45, 46^. We also observed a decrease of ribosomal subunits, including mitochondrial ribosomes, in Pex5 knockdown. Consistent with this, down-regulation of Pex5 and Pex12 had a significant reduction on ribosome biogenesis (FDR < 0.0001, Table 1), suggesting dampened ribosome biogenesis. In addition, proteasome pathway is induced in Pex1 and Pex12 knockdowns (Table 1).

Among all three different Pex knockdowns, protein processing in endoplasmic reticulum pathway is up-regulated (Table 1). Density plot showed that comparing to total gene expression in respective RNAi, Pex5^RNAi^, Pex1^RNAi^ and Pex12^RNAi^ all showed a consistent, albeit modest increase of the expression level involved in ER pathway (Fig. 4A-4C). After examining the pathway closely, we identified nine genes that are commonly induced by all Pex RNAi (Fig. 4D). Some of these genes were not identified in previously venn analysis possibly because they did not reach the cut-off (FC ≥ 1.2, FDR ≤ 0.1). Interestingly, CG30156, CG3061, Ubc7, p47, Hsp70Ba and Hsp70Bb (highlighted in red) all participate in ER-associated degradation (ERAD). Among them, p47, Hsp70Ba and Hsp70Bb are highly induced in all Pex knockdowns, by which Hsp70Bb is induced by more than 6-fold in Pex1^RNAi^. p47 is predicted to have ubiquitin binding activity and its human orthologue can interact with ubiquitinated substrates duWe utilized a previously developedring ERAD and are elevated upon ER stress ^47^. Similarly, several ubiquitin ligase complex components are also up regulated, such as sip3 (HRD1 family) and Der-1. The evidence suggests that there is higher level of unfolded proteins in the ER and induced ERAD activity under peroxisomal stress.

To test whether ER stress is induced under peroxisomal stress, we first examined the level of eIF2◻ phosphorylation. Phosphorylation of eIF2◻ on serine 51 reduces protein translation and diminishes the load of unfolded proteins entering ER ^48^. It was observed that oenocyte-specific *Pex5* knockdown induced eIF2◻ phosphorylation in oenocytes (Fig. 3E-F). As the level of PEX5^C11A^ increased, the level of eIF2◻ phosphorylation was also significantly increased in HEK293 mutant cell line (Fig. 3G-H). Addition of Doxycycline to WT cells did not induce the level of eIF2◻ phosphorylation (Supplementary Fig. 2J). We utilized a previously developed xbp1-EGFP reporter to measure Ire-1-mediated splicing activity ^49^. As reported previously, *xbp1* splicing is enhanced to produce in-frame EGFP in response to ER stress, after dithiothreitol (DTT) treatment (Supplementary Fig. 2K-L). However, the level of spliced *xbp1* was similar in control and *Pex5* knockdown flies (Supplementary Fig. 2K-L). Consistently, the level of spliced XBP1 was similar in PEX5^C11A^ mutant cell line treated with Doxycycline compared with control (Supplementary Fig. 2M). 5’ and 3’ primers were set at the positions 412 and 853 of human XBP1 mRNA, so that the amplified fragment would produce a band at 442 bp and spliced band at 426 bp under thapsigargin-induced ER stress ^50^. Altogether, the data showed that peroxisome import defect increased the expression of ER, ERAD genes and induced ISR independent of *ire-1-xbp1* pathway.

**Figure 3.**
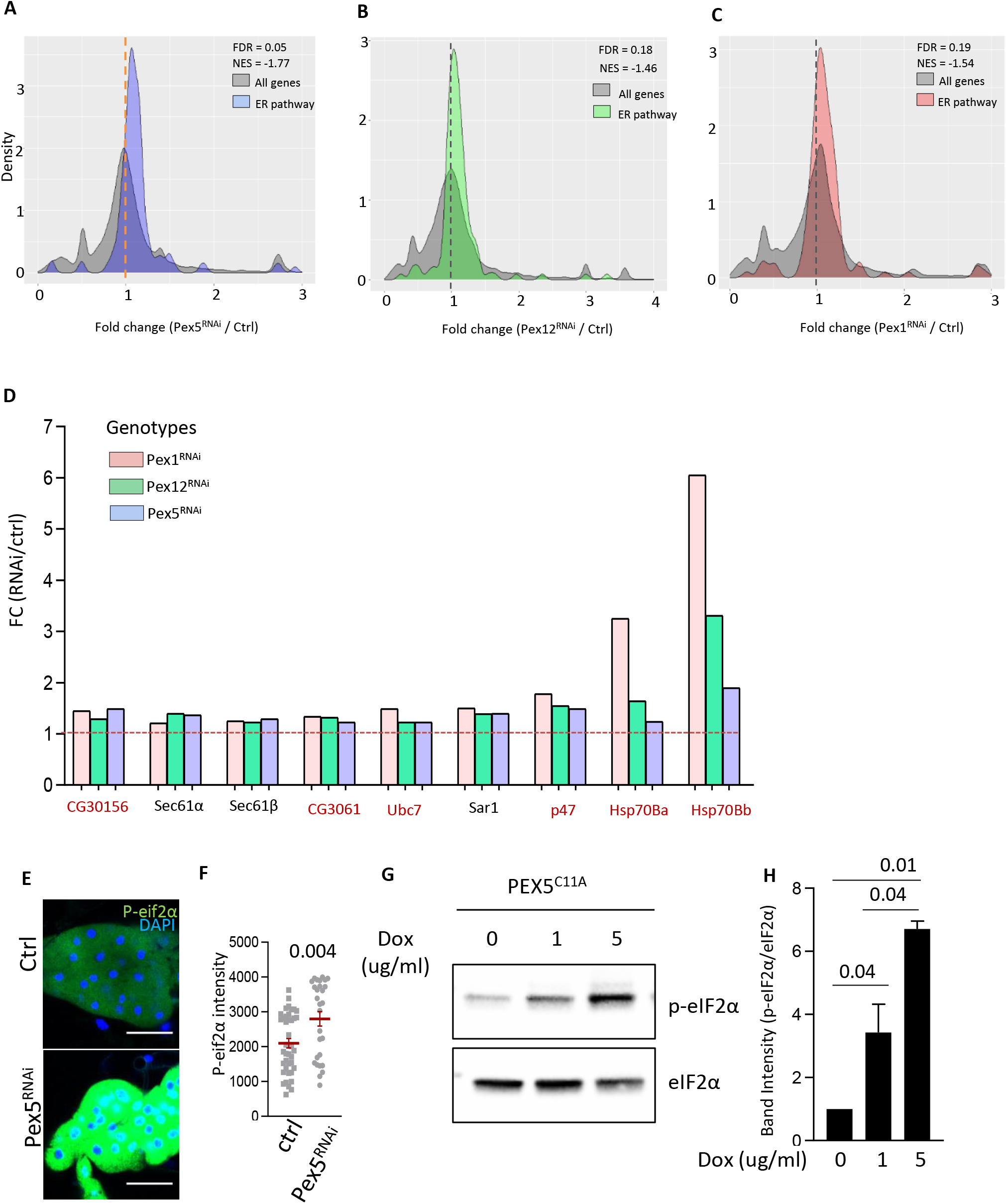
Peroxisomal stress induced endoplasmic reticulum genes and the integrated stress response (ISR). **A-C** Density plot showing the expression fold change (Pex^RNAi^ / Control) of ER pathway genes in oenocytes, in respective Pex^RNAi^. **D** Density plot of fold change in selected genes involved in ERAD pathway across Pex1, Pex12, and Pex5 RNAi samples. **E** Immunostaining of oenocytes with anti-P-eIF2α in control flies (-RU feeding) versus oenocyte-specific *Pex5* RNAi flies (200μM RU). **F** Quantification of P-eIF2α intensity from **E.** N = 6 female flies. **G** Western blot measures P-eIF2α and eIF2α level in PEX5^C11A^ cell line treated with increasing concentration of Doxycycline (0 ug/ml, 1 ug/ml, and 5 ug/ml). **H** Quantification of band intensity ratio (p-eIF2α/eIF2α) in **J**. N = 2-3.

**Figure 4:**
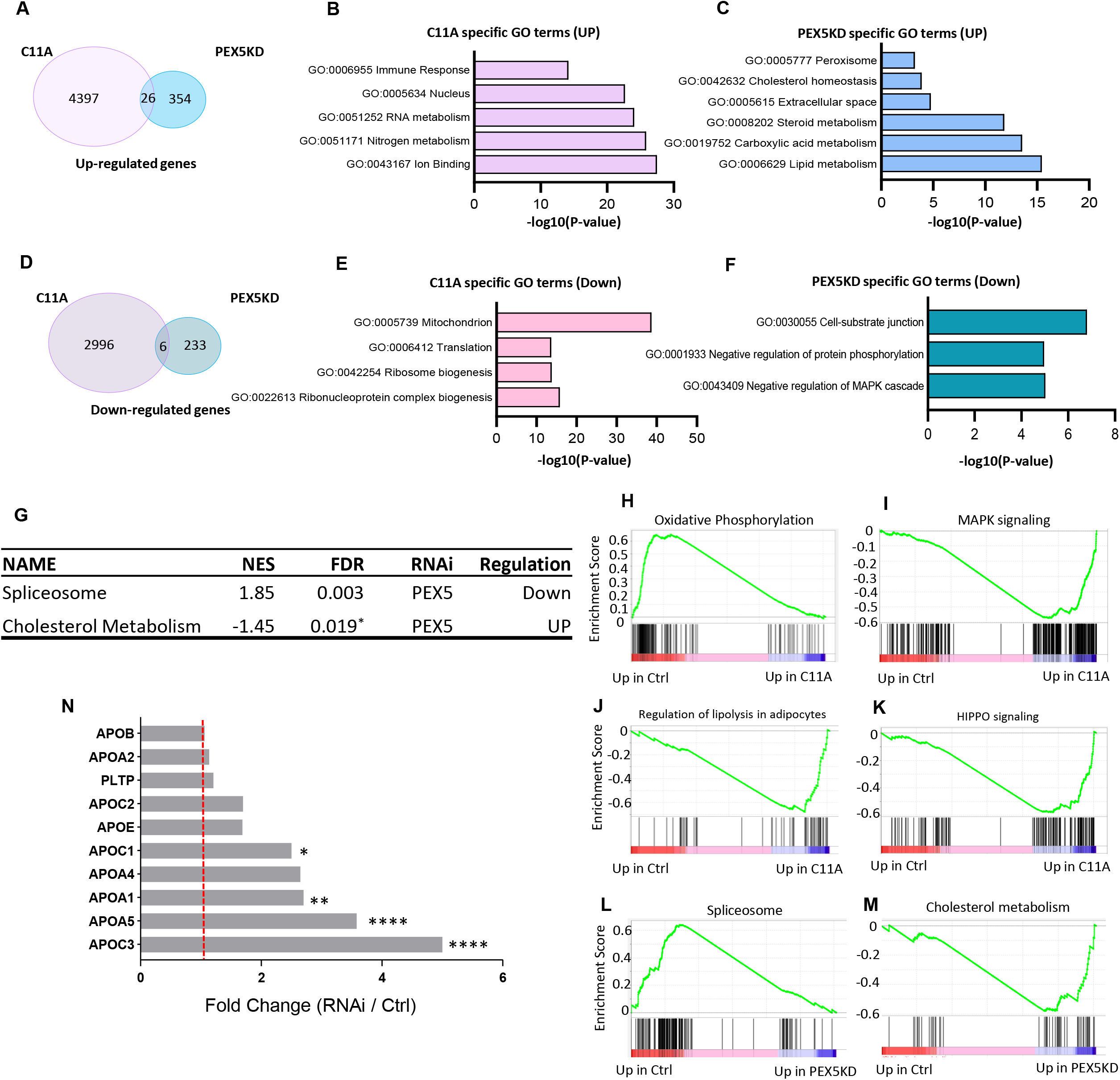
Peroxisome dysfunction up-regulates inflammation and cholesterol pathways in human cells. **A** Venn diagram analysis showing commonly up-regulated genes by PEX5^C11A^ induction and PEX5 RNAi. **B-C** Gene ontology analysis on up-regulated genes only in PEX5^C11A^ or only in PEX5 RNAi cells. **D** Venn diagram analysis showing commonly down-regulated genes by PEX5^C11A^ induction and PEX5 RNAi. **E-F** Gene ontology analysis on down-regulated genes only in PEX5^C11A^ or only in PEX5 RNAi cells. **G** List of enriched pathways from GSEA analysis on PEX5 RNAi cells. * represents P-value. **H-K** GSEA enrichment profiles on PEX5^C11A^: Oxidative phosphorylation, MAPK signaling, regulation of lipolysis in adipocytes, HIPPO signaling. **L-M** GSEA enrichment profiles on PEX5 RNAi: spliceosome, cholesterol metabolism. **N** Selected genes from cholesterol metabolism pathways on PEX5 RNAi cells. Y-axis represents fold change (RNAi / Control).

### Peroxisome dysfunction represses oxidative phosphorylation and induced inflammation, cholesterol metabolic pathway in human cells

To understand how peroxisome stress changes transcriptions in human cell cultures, we performed GO term analysis on DEGs (│fold change│ ≥ 1.2, FDR ≤ 0.1) from PEX5^C11A^ and PEX5-siRNA treated cells. PEX5-siRNA treatment produced 587 DEGs, whereas PEX5^C11A^ elicited dramatic transcriptional response, with 7393 DEGs. Interestingly, these two treatments share few common transcriptional changes: 26 genes commonly induced versus only 6 genes commonly repressed by two treatments (Fig. 3A). Among these 26 commonly up-regulated genes (Supplementary Fig.1G), majority of them regulates cell proliferation (CCNA2, FGFR3, CDCA5, PSMB9 and UBA7) and mitochondrial metabolism (PCK1, ACSM3, ALDH8A1 and BTG2). We examined the specific genes induced by PEX5^C11A^ and PEX5-siRNA. PEX5^C11A^ specifically induced immune response, RNA metabolism, nitrogen metabolism (Fig. 3B), whereas PEX5-siRNA specifically induced peroxisome genes, cholesterol homeostasis, steroid metabolism, carboxylic acid metabolism and lipid metabolism (Fig. 3C). PEX5^C11A^ represses mitochondrial genes, translation, ribosome biogenesis. PEX5-siRNA represses negative regulation of protein phosphorylation and MAPK cascade, indicating an induction of MAPK signaling under PEX5 knockdown, which is consistent with previous findings in oenocytes ^19^.

To further characterize peroxisome-stress induced signaling pathways, we performed GSEA, using a collection of pre-defined gene sets retrieved from KEGG database on *Homo Sapiens*. PEX5^C11A^ significantly regulated 103 signaling pathways, whereas only 2 pathways are significant under PEX5-siRNA (FDR ≤ 0.05). Through GSEA analysis, we found oxidative phosphorylation pathway and valine leucine and isoleucine degradation pathway are downregulated under PEX5^C11A^ treatment (Table 2 and 3G). In oxidative phosphorylation pathway, we found that key components of Complex I-IV of the respiratory chain are down-regulated. For example, NADH dehydrogenase: NDUFV1, NDUFV2, NDUFS2, NDUFS3, NDUFS4, NDUFS6, NDUFS7, NDUFS8 and NDUFSA11, which are components of Complex I and deficiency of which is the most common cause of mitochondrial disease^51^. Cytochrome C1 (CYC1) which is part of the metal centers responsible for electron transfer in complex III, is also downregulated. Almost all cytochrome c oxidase (COX, complex IV) subunits are repressed (COX4l1, COX4l2, COX5A, COX5B, COX6A1, COX6B1, COX6C, COX7A1, COX7A2, COX7A2L, COX7B, COX7C, COX8A and COX15) and many ATPase subunits (ATP6V1G2, ATP6V0C, ATP6V1G1 etc.) are repressed as well. These results suggest that mitochondrial electron transport activity is greatly impaired under peroxisomal stress, which is consistent with previous publications^52–54^. Even though we did not observe decreased oxidative phosphorylation genes in *Drosophila* oenocytes, we observed an altered mitochondria morphology with higher fusion activity under Pex5 knockdown in oenocytes (unpublished observation). This indicates peroxisome and mitochondria are dynamically connected organelles and the response to peroxisome stress is conserved between flies and human.

**Table 2.**
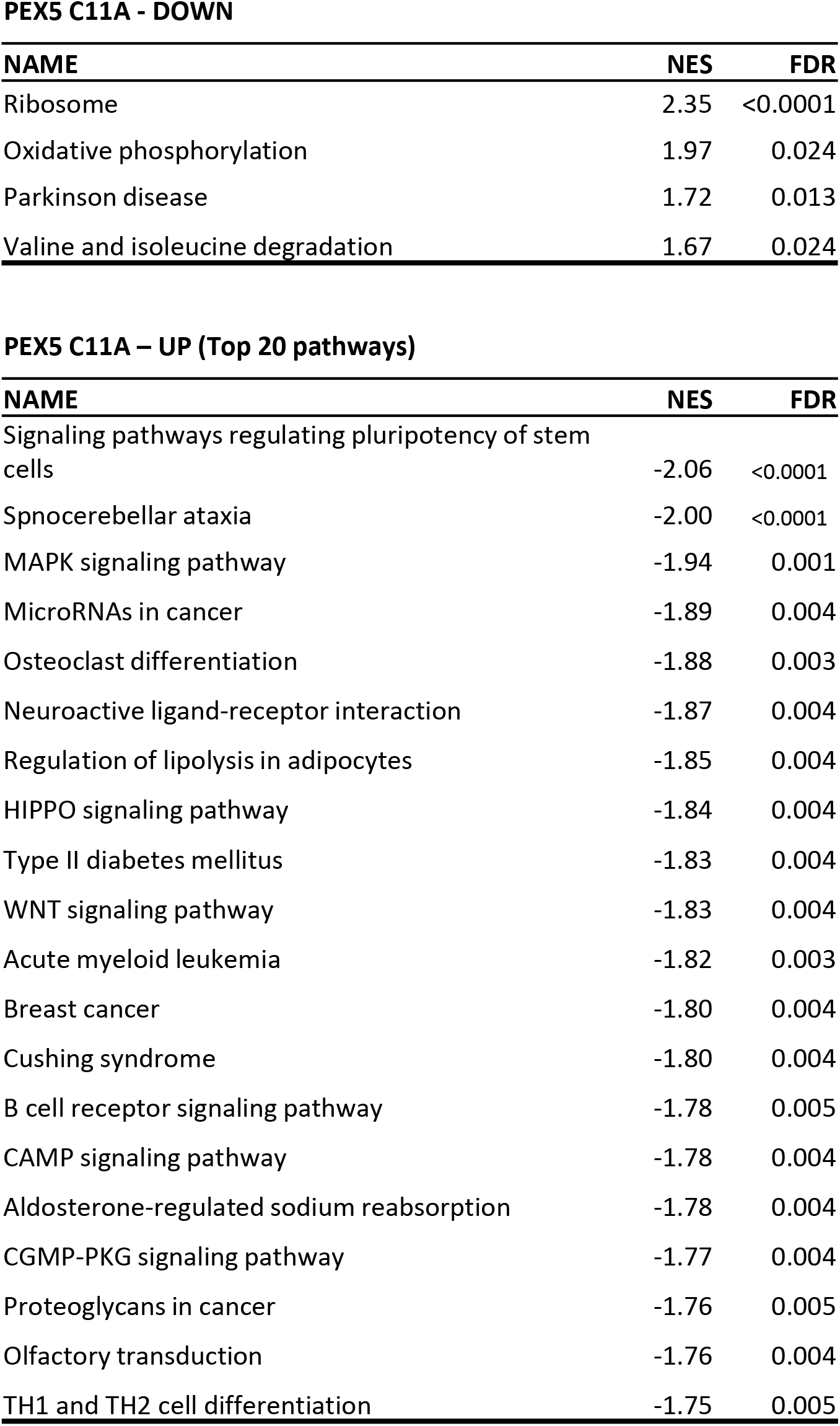

It has been well established that defects in electron transport chain leads to elevated reactive oxygen species (ROS) and can induce inflammatory pathways^55 56^. Interestingly, we also observed activated MAPK signaling, type II diabetes mellitus under PEX5^C11A^ treatment (Table 1). Pex5 knockdown was previously identified to activate MAPK/JNK signaling in *Drosophila 41 Hawthorne St #1* oenocytes ^19^, this suggests a conserved cellular response from peroxisome stress between flies and human. In addition, we also observed an upregulation of HIPPO pathway, including key regulators such as MOB1B, LATS1/2, YAP1. HIPPO pathway which is originally discovered for its regulation on organ size. However, recent evidence has discovered that Hippo signaling is also involved in oncogenesis ^57, 58^ as well as apoptosis ^59–65^. Peroxins mutants exhibit enhanced cell apoptosis and cell death ^66^ (based on unpublished data), our evidence suggest that peroxisome stress induced by PEX5^C11A^ activates apoptosis through HIPPO signaling activation.

PEX5-siRNA treatment produced less significant GSEA pathways comparing to PEX5^C11A^, with only spliceosome showing significant negative correlation with age (NES = 1.85, FDR = 0.03). Spliceosome and RNA processing are also negatively correlated with age in human^67^. which suggests that peroxisome defects are one of the causes for tissue aging. According to GSEA analysis, under PEX5-siRNA induced genes involved in cholesterol metabolism pathway (NES = −1.45, P-value = 0.019) (Fig. 3F). Among the upregulated genes in the pathway, apolipoprotein genes are highly induced, including APOC1, APPOA1, APOA5 and APOC3 (Fig. 3M). APOC3 had a significant 5-fold increase after PEX5 knockdown (Fig. 3M). Apolipoproteins are crucial for lipoprotein metabolism. Not only can guide lipoprotein formation, they also serve as activator or inhibitors of enzymes involved in lipoprotein metabolism ^68^. Notably, single-nucleotide polymorphisms within the apoliprotein (APO) gene cluster, which includes APOA1/C3/A4/5 genes, are strong risk alleles associated with hypertriglyceridemia and increased coronary heart disease (CHD) risk in humans ^69, 70^. APOA5 encodes a protein which is secreted into plasma to control plasma triglyceride (TG) metabolism by acting as an activator of lipoprotein lipase, thus promoting TG catabolism^71^. In contrast, APOC3 encodes a protein that acts as an inhibitor of lipoprotein lipase and high expression of APOC3 have been associated with increased plasma level of TG and hypertriglyceridemia ^72^. Interestingly, APOC3 protein is also a modulator for pro-inflammatory pathways, such as PKCβ and NF-κB in endothelial cells. It is interesting that both APOC3 and APOA5, which have opposing effects on TG, all are induced under PEX5-siRNA. Pex5 perturbation’s effects on plasma TG require further investigations. Many of pathway regulations are cell type or treatment specific. Despite of these differences, we were able to identify several pathways, including MAPK, mitochondrial oxidative phosphorylation, elevated ISR as conserved response to peroxisome dysfunction.

### GSEA uncovers ribosome biogenesis declines as a conserved response in the peroxisomal defects

To identify conserved response to peroxisomal stress, we compared the GSEA results in *Drosophila* oenocytes and in human cells. GSEA showed a conserved downregulation of genes functioning in the KEGG pathway ribosome biogenesis in Pex5^RNAi^, Pex12^RNAi^ (*Drosophila* oenocytes) and PEX5^siRNA^ in human cells (Fig. 5A and Supplementary Fig. 2A). As shown by density plots (Fig. 5A), most of genes in the ribosome biogenesis gene set showed a consistent, although modest, decrease in the expression level between knockdown and control, compared to total gene expression (Fig. 4A-B and Supplementary Fig. 2D-F). A similar trend was observed with ribosome gene set (Fig. 5B), with a decrease in ribosome genes in PEX5^C11A^ and Pex5^RNAi^, including majority of genes encoding mitochondrial ribosomal proteins and ribosomal proteins. Even though RU feeding also induced the ribosome pathway, however the heatmap showed RU and Pex5^RNAi^ showed few similarities on the expression level (Supplementary Fig. 2B).

**Figure 5.**
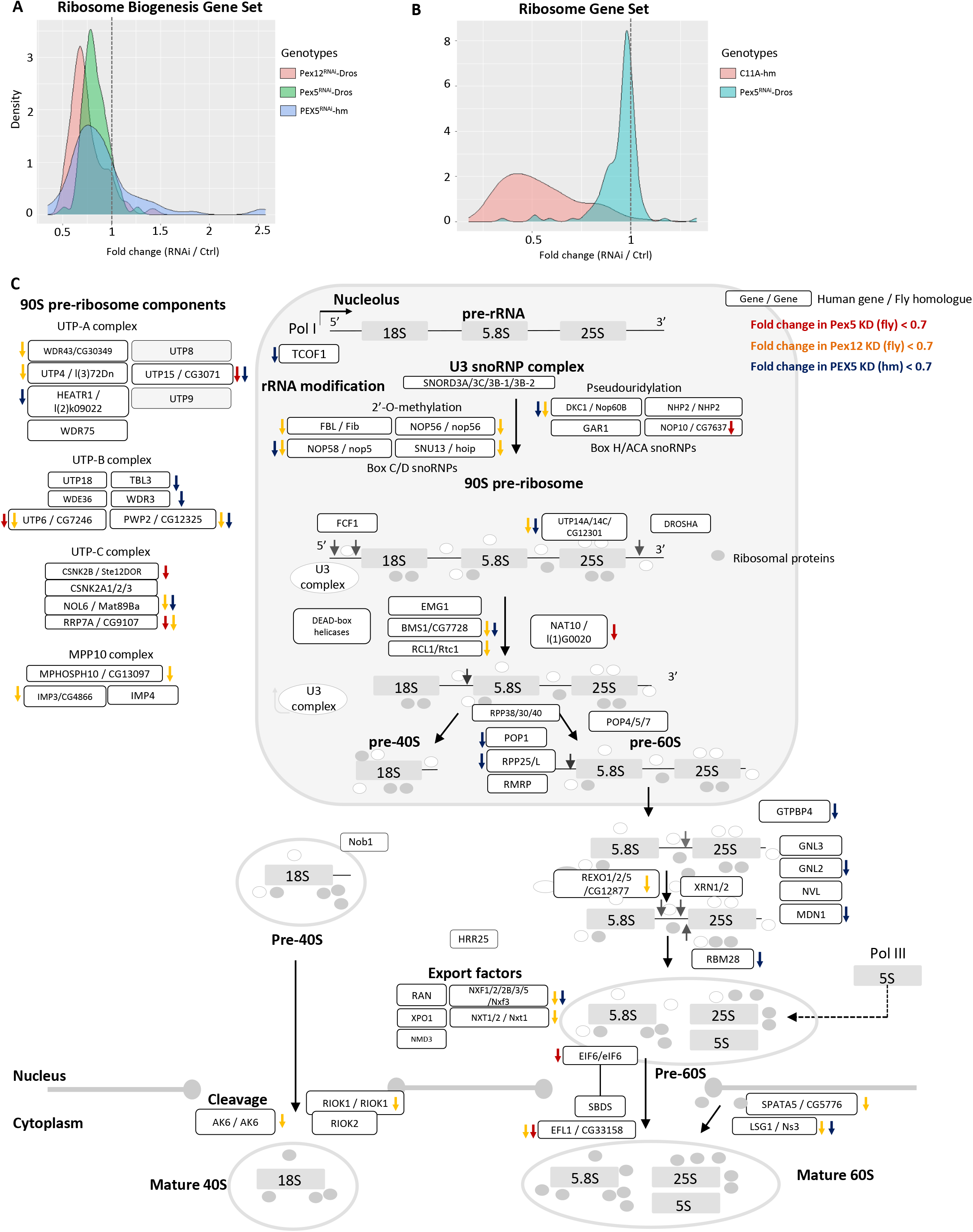
Peroxisome dysfunction represses ribosomal genes in both flies and humans: **A** Density plot showing the fold change (Treatment / Control) of ribosome biogenesis pathway genes in oenocytes and PEX5 RNAi in human cells. **B** Density plot showing the fold change (Treatment / Control) of ribosome pathway genes in oenocytes and PEX5^C11A^ in human cells. **C** Schematic diagram showing ribosome biogenesis pathway, the role of biogenesis factors and their fold change in oenocytes (Pex5, Pex12 RNAi) and human cells (PEX5 RNAi). Genes reduced the expression under treatment for more than 0.7-fold are noted on the diagram (red arrow indicates genes reduced under Pex5 RNAi in oenocytes, yellow indicates Pex12 RNAi, blue indicates PEX5 KD in human cells).

A closer look on ribosome biogenesis pathway revealed that multiple steps regulating the process are repressed. Peroxisomal stress from three different knockdowns all target on 60S ribosome processing and 90S pre-ribosome components (Fig. 5C). The eukaryotes contain two ribosome subunits, the 40S (small) and the 60S (large), which contain four rRNA species (18S, 5.8S, 25S and 5S) and ribosomal proteins (RPs). Ribosome assembly starts in the nucleolus by transcribing a large precursor rRNA (47S pre-rRNA in *Drosophila* and mammals) by RNA polymerase I, from which the mature 18S, 5.8S, and 25S rRNAs are generated ^27^. Our results show multiple steps during ribosome processing are affected under Pex knockdown. For example, NOP58, a core component of the box C/D small nucleolar ribonucleoprotein complex, which controls specific pre-ribosomal RNA processing, is downregulated in PEX5^RNAi^ in human cells. Its fly orthologue, nop5, is also repressed under Pex12^RNAi^ in oenocytes. Ribosome biogenesis protein BMS1 homolog (BMS1), which is crucial for 40S maturation are repressed in human PEX5^RNAi^, same with its orthologue is flies. Additionally, elongation factor like GTPase 1 (EFL1) can assist the release of eukaryotic translation initiation factor 6 (eIF6) release from 60S-ribosome ^73, 74^. Thus, it plays an important role in translational activation. CG33158 is the fly orthologue of EFL1. Interestingly, we found reduced expression level of CG33158 in Pex5^RNAi^ and Pex12^RNAi^ in oenocytes, suggesting the translation activity is dampened under peroxisome stress.

During ribosome biogenesis, the 47S pre-rRNA assembles co-transcriptionally with various *trans-*acting factors and early binding ribosomal proteins of the small subunit. Together a huge macromolecular complex, named 90S pre-ribosome or small-subunit (SSU) processome will form^26, 29, 30^. 90S pre-ribosome is composed of 60-70 non-ribosomal factors, with most being called U three proteins (Utp) ^26^, and they can form structurally autonomous subcomplexes (UTP-A, UTP-B, UTP-C, Mpp10-Imp3-Imp4, U3 snoRNP, and Bms1-Rcl1 modules). These modules will associate with pre-rRNA in a sequential and hierarchical order ^75–77^. UTP-A and UTP-B are the earliest 90S modules that assemble on the 5’-external transcribed spacer (ETS) of the pre-rRNA, whereas UTP-C and other complexes bind later. It has been reported that UTP-A complex is critical for the initial cleavage of 5’ETS site on 47S pre-rRNA, thus regulating the ribosome biogenesis ^78^. We found multiple genes in SSU processome are downregulated under peroxisome stress. UTP15 small subunit processome component (UTP15), nucleolar protein 6 (NOL6), PWP2 small subunit processome (PWP2), are all repressed in PEX5^RNAi^ in human cells or their fly orthologues are repressed in oenocytes. Because UTP-A, UTP-B and UTP-C components are largely dampened under peroxisomal stress, we sought to verify whether peroxisomal stress also caused pre-rRNA processing, especially at the 5’-ETS removal steps.

pre-rRNA processing starts with the removal of the 5’ and 3’-ETS, and the internal transcribed spacers 1 and 2 (ITS1 and ITS2, respectively) (Fig. 6A). To test whether pre-rRNA processing is impaired under peroxisomal stress, we designed primers against the 5’-ETS region and 3’-ETS regions in Pex5^RNAi^ oenocytes and PEX5^C11A^ induced human cells (Fig 6A). We discovered strong accumulation of higher level of 5’-ETS and ITS1 could indicate a defect in rRNA processing, or induced level of rRNA transcription (Fig. 6B-6C). Next, we measured uncleaved ETS using primer 1 and 2 as shown in figure 6A, normalized to 18S (primer 3 and 4) which represents total level of rRNA transcribed ^79^. Higher relative expression value is indicative of accumulated unprocessed rRNA. Pex5 knockdown specifically in oenocytes can increase the fraction of 5’ETS-18S unprocessed rRNA (Fig. 6D), without significantly affecting the total rRNA level (Fig.6E). We also found a consistent increase of unprocessed rRNA in human cells after PEX5^C11A^ induction, but not 18S level (Fig. 6G-6H). To further verify whether ribosome synthesis is impaired, we measured the localization of ribosomal protein S6 (Rps6) under peroxisomal stress. The rationale for this assay is based on the observation that defects if biogenesis pathway can cause ribosomal subunit export defects ^80, 81^. Rps6 is an early assembling 40S subunit ribosomal protein ^82–84^, allowing us to score early defects in 40S synthesis. Indeed, we found an accumulation of RPS6 in the nucleus in Pex5^RNAi^ oenocytes, after normalized to total intensity (Fig. 6I – 6J). We also identified higher percentage of cells contain nucleus rpS6 signal in PEX5^C11A^ expressing human cells, comparing to control (Fig 6K-6L). These results indicate a defect in rRNA processing, instead of increased rRNA transcription, under peroxisome disruption.

**Figure 6.**
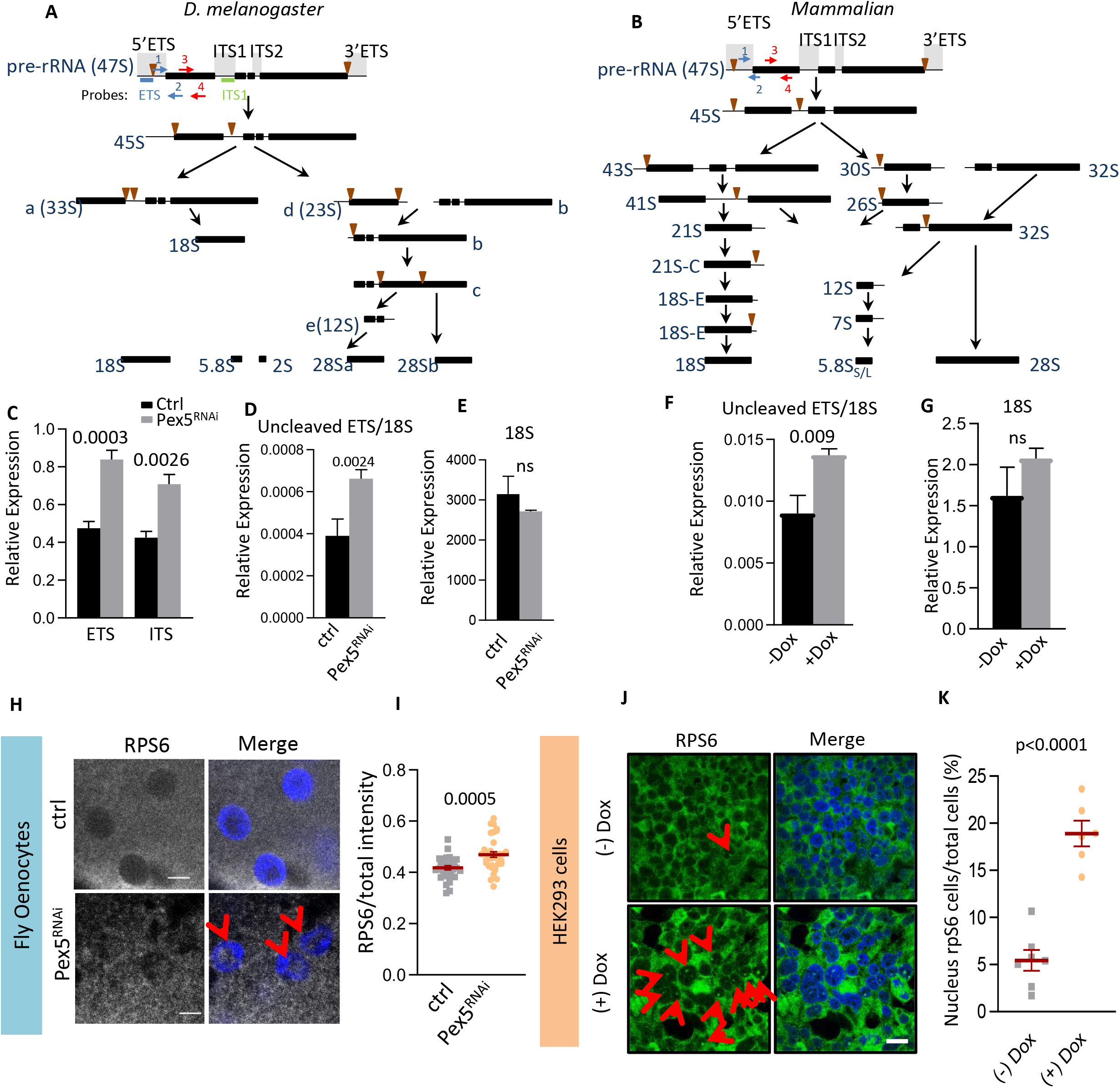
Impaired ribosome biogenesis genes in human and flies: **A-B** Overview of *D. melanogaster*, and Mammalian rRNA biogenesis Pathways. rRNA biogenesis intermediates are generally conserved among vertebrates; 5.8S rRNA is cleaved into 2S and a short 5.8S, and 28S rRNAs are cleaved into 28Sa and 28Sb. Blue arrows indicate the locations of the primers for ETS. Green label indicates primer locations for ITS1. Red arrow indicates primers for 18S. **C-F** The relative amounts of the ETS, ITS1, unprocessed ETS/18S ratio, 18S were measured by qPCR using RNA isolated from control oenocytes or Pex5 RNAi oenocytes. Data are presented as the mean ± s.e.m (n = 2). **G-H** The relative amounts of the unprocessed ETS/18S ratio, 18S were measured by qPCR using RNA isolated from control cells (-DOX) or PEX5^C11A^ (+DOX) cells. Data are presented as the mean ± s.e.m (n = 3). **I-L** Analysis of ribosome proteins (RPs) nucleolar localization. **I** Depletion of Pex5 in oenocytes results in the accumulation of RpS6 nucleolus. Scale bar shows 3.3μm. **J** Quantification of RpS6 intensity in nucleolus normalized to total intensity within the cell. **K** Expressing PEX^C11A^ (+Dox) results in the accumulation of RpS6 nucleolus in HEK293 cells (n=6). Scale bar shows 20μm. **L** Quantification of number of cells containing RpS6 signal in nucleolus normalized to total number of cells, in control (-Dox) and PEX5^C11A^ expressing cells (+Dox) (n = 6).

## Discussion

The peroxisome is the important metabolic site to perform β-oxidation of fatty acids and the degradation of toxic hydrogen peroxide, yet it has received little respect until now. Although peroxisomal dysfunction has been linked to severe diseases in man and aging ^15, 17, 19, 85^, whether cells can mount defensive mechanism and how cells respond to peroxisome stress remain to be determined. Here we characterized a variety of transcriptional alterations under multiple peroxisomal stress in both *Drosophila* oenocytes and human cell cultures. We show that defects in peroxisome import process can elicits distinct transcriptional response in *Drosophila* oenocytes. Peroxisomes are highly interconnected with other organelles, such as ER and mitochondria, so that peroxisome dysfunction can induce gene expression targeting to other organelles. Specifically, peroxisome defects in oenocytes up-regulates genes involved in ER and ERAD. PEX5^C11A^ induction induced genes involved in oxidative phosphorylation and altered mitochondrial ribosome genes’ expression. PEX5^RNAi^ specifically up-regulated genes involved in cholesterol metabolism and transport. We also identified conserved decrease in ribosome biogenesis and ribosome between *Drosophila* and human cells, highlighting it as a direct and crucial mechanism to peroxisome stress.

### Different importing peroxins control distinct transcriptional processes

Maintaining peroxisomal activity is a highly dynamic and complicated process, facilitated by peroxisomal biogenesis factors (peroxins). In fact, most of the peroxins identified so far are known to be directly involved in different stages of peroxisomal matrix protein import ^86^. Newly synthesized and folded peroxisome proteins from cytosol are targeted to the organelle in a post-translational manner^86^. The protein import process into peroxisome is involved with multiple steps, including cargo recognition, docking of the cargo/receptor-complex at the peroxisome membrane, cargo translocation, cargo release into the peroxisomal matrix and receptor recycling (Fig. 1A). Disrupting key regulators during these processes can compromise the peroxisome import activity and lead to severe physiological consequences^19, 35^, including development of PBDs. Even though these Peroxin genes work cooperatively to maintain peroxisome activity, it remains unknown whether their sole activity is to regulate peroxisome biogenesis.

Interestingly, our transcriptomic analysis on multiple import-regulatory genes (*Pex5, Pex12* and *Pex1*) suggests that they may regulate other cellular processes. First, we found that *Pex1* and *Pex5* RNAi produced high level of transcriptional changes comparing to *Pex12* (1055 and 1056 DEGs versus 248 DEGs). This might be because the severity of peroxisome import is different, because *Pex12* dsRNA in S2 human cells show less diffusion of cytosolic GFP-PTS comparing to *Pex1* and *Pex5 ^35^.* However, both our GO ontology and GSEA analysis suggest that even *Pex1* and *Pex5* knockdown induced different genetic pathways. *Pex5* knock down induced GO terms related to oxidation-reduction and glutathione pathways, whereas *Pex1* dramatically induced genes involved in translation, ribosome, and proteasome (Table 1). The discrepancies in their transcriptomic profiles, despite sharing the same phenotype in peroxisome import, suggest that they regulate different processes other than Pex5 recycling. Pex1 and Pex6 both belong to the AAA ATPase family, which is known to be involved in protein quality control, including folding, assembly, transport, and protein degradation ^87^. This could explain the induction of translation and proteasome in *Pex1^RNAi^.* Pex1, Pex15 and Pex6 will form an exportomer and deficiency of these components can trigger pexophagy in mammalian cells ^88^. Another study found that a missense mutation of Pex1 can ameliorated the defects of Pex6 mutant without restoring PEX5 recycling, possibly through prevention of pexophagy, suggesting other functions Pex1-Pex6 complex ^89^. In support of this view, studies have unexpectedly identified *PEX6* regulating mitochondrial inheritance in daughter cells in yeast. Seo et al. showed that PEX6 overexpression was able to rescue the deficit in Atp2p (the β-subunit of mitochondrial F_1_, F_0_-ATPase ^90^) mutant by improving its import into mitochondria. Studies suggest that Pex1 was able to translocate between mitochondria and peroxisomes^91^, where they showed that mitochondria localized PEX6-PEX1-PEX26 complex can rescue PEX26 deficient cells. Altogether, our findings provide new insights in peroxisome biogenesis disorders, peroxins functions and their significance of other organelles’ activities.

### Peroxisome and inter-organelle communications

To optimize their activities, peroxisomes must coordinate with other organelles frequently. No peroxisome is in isolation. A growing body of evidence showed that peroxisome share physical membrane contact sites with other organelles (see an excellent review here ^4^). This suggests there is intimate interaction between peroxisomes with these organelles, which is important for controlling processes such as metabolism and organelle proliferation. In agreement of this concept, we identified changes of gene expression for other organelles, especially for ER and mitochondria, under various Pex knockdowns.

Our results of PEX5^RNAi^ in HepG show significant up-regulation of cholesterol metabolism (Fig. 3C and 3F). Previous study shows that cholesterol accumulates drastically in animals and human patients with peroxisomal disorders, as a result of disrupted peroxisome-lysosome contact sites ^92^. The majority of cellular cholesterol (60% - 80%) is localized at the plasma membrane (PM) ^93^. Cholesterol is synthesized in the ER or it can be imported exogenously and stored in the lysosome. Cholesterol can be transported from the ER and lysosome to the PM. Recently it was shown that peroxisome mediates the transfer of cholesterol from lysosome to PM. A peroxisome-lysosome contact site was shown to be required for this transport ^92^. An up-regulation of cholesterol metabolism genes could be a compensatory mechanism to degrade excess cholesterol accumulated inside the cells.

Besides lysosome, peroxisomes also actively interact with ER and mitochondria. Our results show that mitochondrial ribosomal subunits are repressed under Pex5 knockdown in oenocytes (Table 1), as well as in PEX5^C11A^ mutant (Table 2). Consistently, genes involved in oxidative phosphorylation are largely repressed in PEX5^C11A^, which was also been observed in Pex16 KO ^94^. Our results suggest that mitochondria function is altered under peroxisome stress. In fact, previous studies as well as our observation have found that mitochondria underwent functional and morphological changes under Pex5 and Pex16 deficiencies ^94, 95^. In these studies, they consistently observed a more fused mitochondria structure, reduced level of the respiratory chain activities. Park et. al identified that peroxisome-derived phospholipids are responsible peroxisome dysfunction induced mitochondrial abnormalities ^94^. Park et. al inhibited plasmalogen synthesis by knocking down expression of GNPAT, an enzyme responsible for the first step in plasmalogen production. GNPAT knockdown was able to recapitulate phenotypes with Pex16 knockdown, including altered mitochondrial morphology and decreased mtDNA. Interestingly, they also found dietary supplementation of plasmalogen precursors can rescue thermogenesis in Pex16-KO mice. However, whether plasmalogen level is the sole contributor to mitochondrial dysfunction and why there are tissue specific mitochondria abnormalities under peroxisome deficiency, these questions still require investigation.

Our results suggest different Pex knockdowns commonly induce genes VLCFA synthesis, whose protein products can localize on ER (e.g. *spidey* and *Cyp6g1*). Biosynthesis of VLCFA takes place at the ER and can be metabolized in peroxisomes ^96, 97^. Our findings suggest two organelles work closely to regulate metabolism of VLCFA. EM images showed that two organelles are adjacent to each other and that ER membrane can wrap around peroxisomes ^98–101^. In fact, peroxisome-ER crosstalk is important for many lipid-related metabolic pathways, including the biosynthesis of ether-phospholipids, production of polyunsaturated fatty acids, bile acids, isoprenoids, and cholesterol.

Interestingly, we found peroxisome dysfunction in oenocyte cells can trigger protein process pathway in the ER and ERAD (including CG30156, CG30161, Ubc7, p47, Hsp70Ba and Hsp70Bb), which is possibly due to elevated ER stress (Fig. 3). Consistently, it has been previously identified that peroxisome-deficient *Pex2* and *Pex5* knockout mice had elevated level of ERAD gene expression, ER stress and ER abnormalities, prominently through PERK activation ^95, 102^. How does peroxisome dysfunction lead to the ER stress? Possible reasons include perturbed flux of mevalonate metabolites, changes in fatty acid levels or composition and increased oxidative stress ^103^. Specific induction of PERK under peroxisome stress, not activation of Xbp1 splicing, suggesting peroxisome stress induced a specific cellular response distinct from protein misfolding induced ER stress. Genetic experiments and biochemical assays should be utilized to discern the nature of peroxisome induced ER stress.

### Peroxisome in ribosome biogenesis and protein homeostasis

Our results identified decreased transcription on ribosomal proteins and ribosome biogenesis factors, as a conserved response to peroxisome stress between humans and *Drosophila*. This is also the first study to verify the reduced ribosome biogenesis under Pex5^RNAi^ in oenocytes and PEX5^C11A^ mutant in human cells. Our results show that the 5’ cleavage process is inhibited under Pex5 knockdown. Our result indicates that peroxisome defects may specifically regulate ribosome biogenesis by altering ribosome biogenesis process, which might be more efficient in dealing with transient stress. In addition, we also observed an accumulation of RPS6 in the nucleus under Pex5 stress, which is often an indicator of ribosome biogenesis defects, specifically involved with 40S maturation process ^80^. However, it is highly possible that 60S maturation is also affected, judged by down-regulated expression of several crucial genes: Midasin (MDN1)^104^, GTPBP4 ^105, 106^ and EFL1^73, 74^. Overall, our results indicate a down-regulation of ribosome availabilities, and possibly lower translational activities under peroxisome stress.

Why multiple peroxisome defects converge to impair ribosome biogenesis? It is possible that nucleolus is sensitive to peroxisomal or mitochondrial ROS level, which can hamper the function of ribosome biogenesis proteins that primarily localize in the nucleolus. However, very little is known on what stresses can alter nucleolus protein activity or composition. Further research is needed to investigate this area.

It has been well established that reduced ribosome proteins and ribosome biogenesis factors can increase longevity in model organisms (see ^23^ for review). The evidence also suggest that reduced ribosome biogenesis is a protective mechanism to confer peroxisomal stresses. Being a regulator of ribosome biogenesis, Pex5’s effect on lifespan had been controversial. In Δpex5 yeast cells, there was a strong reduction in chronological lifespan^20^. However, post developmental knockdown of prx-5, a *C. elegans* homolog for mouse *Pex5*, increased the worm’s lifespan ^21, 22^. The paradoxically increased lifespan by peroxisome disruption could be attributed to decreased level of ribosome and translational activities. It will be interesting to see whether post developmental knockdown of Pex5 in oenocytes can also prolong lifespan through the regulation on ribosome biogenesis. In addition, how peroxisome import defect reduces ribosome biogenesis will be an interesting avenue to investigate.

## Methods

### Plasmid Construction

Human Pex5 cDNA from the Mammalian Gene Collection (MGC) was purchased from Dharmacon. The hPEX5 was amplified by PCR using the forward and reverse primers (5’-CACTATAGGGAGACCCAAGCTTATCTAGACATGGCAATGCGGGAGCT-3’ and 5’-TCTTACTTGTCATCGTCGTCCTTGTAGTCGCCCTGGGGCAGGCC-3’) and introduced between XhoI and BamHI sites in c-Flag pcDNA3 (addgene #20011) to generate Flag-tagged hPEX5 using NEBuilder^®^ HiFi DNA assembly Master mix (New England Biolabs). Site-directed mutagenesis for amino acid substitution was performed using the Q5^®^ Site-directed mutagenesis kit (New England Biolabs) according to the manufacture’s instruction. The primers for C11A mutant were 5’-GGAGGCCGAAgctGGGGGTGCCAACC-3’ and 5’-ACCAGCTCCCGCATTGCC-3’.

To generate Tetracycline inducible PEX5C11A plasmid, we modified pMK243 (Tet-OsTIR1-PURO) from Masato Kanemaki (Addgene #72835). pMK243 was digested by BgIII and MluI to remove OsTIR sequence. Flag-PEX5C11A was amplified by PCR using the forward and reverse primers (5’-gattatgatcctctagacatatgctgcagattacttgtcatcgtcgtccttgtagt-3’ and 5’-tcctaccctcgtaaagaattcgcggccgcaatggcaatgcgggagctggt-3’) and introduced between BgIII and MluI sites in digested pMK243 plasmid to generate Tet-PEX5C11A-PURO plasmid. All plasmids are confirmed by Sanger sequencing.

### Generation of CRISPR Knock-in HEK293 cells expressing PEX5C11A

HEK293 cells were maintained in Dulbecco’s Modified Eagle Medium containing 10% fetal bovine serum (FBS), with penicillin and streptomycin. To generate stable cell line, we followed the protocol described in Natsume et al. 1×10^6 HEK293 cells were plated in one well of a 6-well plate. After 24hrs, 800ng of AAVS1 T2 CRISPR in pX330 (Addgene #72833) and 1ug of Tet-PEX5C11A-PURO were transfected using Effectene (Qiagen) according to the manufacturer’s instructions. After 48hrs, the cells were detached and diluted at 10 to 100 times in 10ml of medium containing 1ug/ml of puromycin and transferred to 10 cm dishes. The selection medium was exchanged every 3 to 4 days. After 8 to 10 days, colonies were marked using a marker pen under a microscope, picked by pipetting with 10ul of trypsin-EDTA, and subsequently transferred to a 96-well plate. 100ul of the selection medium was added. When the cells were confluent, they were transferred to a 24-well plate. After 2-3 days, the cells were transferred to a 6-well plate. After 2-3 days, cells were detached and half of the cells were frozen, and the rest were used for genomic DNA preparation.

### Genomic DNA isolation and PCR

To prepare genomic DNA, cells were lysed in buffer A solution (100mM Tris-HCl [pH7.5], 100mM EDTA, 100mM NaCl, 0.5% SDS) followed by incubation at 65°C for 30min. Buffer B (1.43M potassium acetate, 4.28M lithium chloride) was added and incubated on ice for 10min. After centrifuge at 12,000 rpm for 15 min, the supernatant was transferred to new microtube and added isopropanol. After isopropanol precipitation, DNA pellets were washed in 70% ethanol and resuspended with DNAse-free Water.

To verify Tet-PEX5C11A-PURO insertion into AAVS1 locus, genomic PCR was performed using Q5^®^ High-Fidelity DNA polymerase (New England BioLabs) according to the manufacturer’s instruction. 5’-cgtttcttaggatggccttc-3’ and 5’-agaaggatggagaaagagaa-3’ were used for WT cell validation. 5’-cgtttcttaggatggccttc-3’ and 5’-ccgggtaaatctccagagga-3’ were used for Tet-PEX5C11A-PURO integration.

To test Xbp1 splicing activity, control cells (-DOX) or PEX5^C11A^ expressing (+DOX) cells’ total RNA was prepared and amplified by RT-PCR to detect spliced XBP1 mRNA ^50^ (5’-CCTTGTAGTTGAGAACCAGG-3’ and 5’-GGGGCTTGGTATATATGTGG-3’) or GAPDH mRNA (5’-accatcttccaggagcgaga-3’ and 5’-gggccatccacagtcttctg-3’). Resulting products were subjected to 2% agarose gel electrophoresis.

### Fly husbandry and strains

The following RNAi lines were used in the KD experiments: y[1] v[1]; P{y[+t7.7] v[+t1.8]=TRiP.HMJ21920}attP40 (BDSC # 58064, Pex5 RNAi), y[1] sc[*] v[1] sev[21]; P{y[+t7.7] v[+t1.8]=TRiP.HMC03536}attP2/TM3, Sb[1] (BDSC # 53308), y[1] v[1]; P{y[+t7.7] v[+t1.8]=TRiP.HM05190}attP2 (BDSC # 28979). The control line used for the KD experiments is yw, a gift from Rochele Lab. All fly lines are crossed to oenocyte specific gene-switch driver (yw; PromE800-GS-gal4/Cyo;+), a gift from Heinrich Jasper.

Female flies were used in all experiments. Flies were maintained at 25°C, 60% relative humidity, and 12-h light/dark cycle. Adults and larvae were reared on a standard cornmeal and yeast-based diet, unless otherwise noted. The standard cornmeal diet consists of the following materials: 0.8% cornmeal, 10% sugar, and 2.5% yeast. RU486 (mifepristone, Fisher Scientific) was dissolved in 95% ethanol, and added to standard food at a final concentration of 100◻μM for all the experiments. Gene KD or overexpression was achieved by feeding flies on RU486 food for 5-6 days, before RNA isolation.

### Fly oenocyte RNA isolation

Adult tissues oenocytes (20 females per replicate) were all dissected in cold 1◻×◻PBS before RNA extraction. For oenocyte dissection, we first removed FB through liposuction and then detached oenocytes from the cuticle using a small glass needle. Tissue lysis, RNA extraction was performed using RNeasy Micro kit from QIAGEN (catalog number 74034) with the following modifications. Samples were first collected in 1.7ml centrifuge tubes. Dissected tissues were saved in 150ul of Buffer RLT (a component of RNeasy kits) with 143mM β-mercaptoethanol on ice during the dissection. Samples were then put at room temperature for 3 minutes prior to lysis and centrifuged at 7.5xg for 3minutes. Additional 150ul buffer RLT was added and pellet pestle grinder (Kimble pellet pestles, catalog number 749540-0000) was used to lyse the tissue for 1 minutes. 20ng carrier RNA was added to the cell lysates. Freezed samples was thawed in 37°C water bath for 1 minute. Total RNA was extracted using RNeasy Plus Micro columns following company manual.

### Drosophila oenocyte RNA-seq library construction and sequencing

RNA-seq libraries were constructed using 100 ng of total RNA and NEBNext Ultra II RNA Lib Prep (NEB, Ipswich, MA, USA. Catalog number: E7770L). RNA concentrations were measured using Qubit RNA BR Assay Kit (Thermo Fisher Scientific, catalog number: 10210). Poly(A) mRNA was isolated using NEBNext Oligo d(T)25 beads and fragmented into 200_Jnt in size. Purification of the ligation products are performed using Beckman Coulter AMPURE XP (BECKMAN COULTER, catalog number: A63880). After cDNA synthesis, each cDNA library was ligated with a NEBNext adaptor and barcoded with an adaptor-specific index (NEBNext^®^ Multiplex Oligos for Illumina, NEB, catalog number: E7335S). Twelve libraries were pooled in equal concentration and sequenced using Illumina HiSeq 3000 platform (single end, 150bp reads format).

### Human cells library preparation

HEK293 expressing Tet-PEX5C11A-Puro cells were cultured in Dulbecco’s Modified Eagle Medium containing 10% fetal bovine serum (FBS), with penicillin and streptomycin. HepG2 cells were cultured in Minimum Essential Medium containing 10% FBS, 1mM sodium pyruvate, 1X MEM Non-Essential Amino Acids solution (ThermoFisher, #11140050) with penicillin and streptomycin. Cells were incubated in a 37°C incubator in an atmosphere of 5% CO2 in air.

To prepare RNA-seq libraries, 4×10^5 HEK293 expressing Tet-PEX5C11A-Puro cells were seeded in a 12 well plate. After 1 day, 1ug/ml of doxycycline was added to the cells to induce PEX5C11A in the cells. After 72hrs, total RNA was isolated from the cells with or without doxycycline treatment. 3×10^5 HepG2 cells were seeded in a 6-well plate. Transfection of siRNA into HepG2 cells was performed with lipofectamine RNAiMAX (ThermoFisher) according to the manufacturer’s instructions. On-TARGET plus human PEX5 siRNA (Dharmacon # L-015788-00-0005) was used for knockdown of PEX5 in HepG2 cells. For negative control, On-TARGET plus non-targeting siRNA (Dharmacon # D-001810-02-05) was used. After 72hrs, total RNA was isolated from the cells. All samples were treated TURBOTM DNase (Thermofisher #AM1907) to remove all traces of DNA. RNA-seq libraries were constructed using 1ug DNA-free total RNA with NEBNext Ultra II RNA Library Prep Kit for Illumina (New England Biolabs, Ipswich, MA, USA. NEB#E7770). Poly (A) mRNA was isolated using NEBNext Poly(A) mRNA Magnetic isolation module (New England Biolabs, Ipswich, MA, USA. NEB#E7490). After first strand and second strand and second strand cDNA synthesis, each cDNA library was ligated with a NEBNext adaptor and barcoded with an adaptor-specific index. Twelve libraries were pooled in equal concentrations and sequenced using Illumina HiSeq 3000 platform.

### RNA-Seq data processing and differential expression analysis

FaastQC (v0.11.8) was first performed to check the sequencing read quality. Then sequence alignment and mapping was performed using the Spliced Transcripts Alignment to a Reference (STAR) software (v2.7.3a) ^107^. The raw reads were mapped to *D. melanogaster* genome (BDGP Release 6) or Genome Reference Consortium Human Build 38 (GRCh38). Reads mapped were then counted with summarizedOverlaps function using “Union” mode in R. Counts are then analyzed in DESeq2 (v1.26.0) ^108^ for batch control analysis and test for differential expression.

### Principal component analysis (PCA), heatmap and expression correlation plot

PCA graph was generated using plotPCA function of R package DESeq2^108^. Heatmaps and hierarchy clustering analysis were generated using heatmap.2 function of R package gplots. The density plots were plotted using R package ggplot.

### Gene set enrichment analysis (GSEA)

For GSEA analysis, all pre-defined set of 132 KEGG pathways in Drosophila were downloaded from KEGG. Text was trimmed and organized using Java script. Normalized counts were used as input for parametric analysis and organized as suggested by GSEA tutorial site (GSEA ^109, 110^). Collapse dataset to gene symbols was set to false. Permutation type was set to gene set; number of permutations was set to 1000; enrichment statistic used as weighted analysis; metric for ranking genes was set to Signal to Noise.

### Gene ontology and pathway analysis

Functional annotation analysis of differentially expressed genes was performed using STRING^111^ or DAVID^112^. GO terms (Biological Process, Molecular Function, Cellular Component), KEGG pathway, INTERPRO Protein Domains and Features, were retrieved from the analysis.

### Quantitative real-time polymerase chain reaction (qRT-PCR)

qRT-PCR was performed using Quantstudio 3 Real-Time PCR system and SYBR green master mixture (Thermo Fisher Scientific, USA Catalog number: A25778). All gene expression levels were normalized to Rpl32 (in *Drosophila*), GAPDH (in humans) by the method of comparative Ct^113^. Mean and standard errors for each gene were obtained from the averages of three biological replicates, with two technical repeats.

### Western Blotting

5×105 HEK293-PEX5C11A cells were seeded in 6-well plates. After one day, cells were treated with or without doxycycline for 3 days. The proteins were extracted in NP-40 cell lysis buffer (Thermo Fisher Scientific, #FNN0021) with HaltTM phosphatase inhibitor Cocktail (Thermo Fisher Scientific, #78420). Protein samples were denatured with Laemmli sample buffer (Bio-Rad, Cat# 161-0737) at 95°C for 5 min. Then proteins were separated by Mini-PROTEAN^®^ TGX Precast Gels (Bio-Rad). Following incubation with primary and secondary antibodies, the blots were visualized with Pierce ECL Western Blotting Substrate (Thermo Scientific). The following antibodies were used: anti-phospho eIF2α (Cell Signaling Technology, #9721, 1:2000), anti-eIF2α (Cell Signaling Technology, #5324, 1:3000).

### Immunostaining

Anti-RpS6 (Cell Signaling, catalog number 2317) at concentration of 1:100 was used to stain oenocytes’ RpS6^114^. Anti-RpS6 at concentration of 1:50 was used to stain human cells. Anti-phospho-eIF2α (Ser 51, Cell Signaling, catalog number 9721) at concentration of 1:100 was used. Secondary antibodies were obtained from Jackson ImmunoResearch.

Adult oenocyte tissues were dissected in PBS and fixed in 4% paraformaldehyde for 15◻min at room temperature. Tissues were washed with 1x PBS with 0.3% Triton X-100 (PBST) for three times (~5◻min each time), and blocked in PBST with 5% normal goat serum for 30◻min. Tissues were then incubated overnight at 4◻°C with primary antibodies diluted in PBST, followed by the incubation with secondary antibodies for 1◻h at room temperature. After washes, tissues were mounted using ProLong Gold antifade reagent (Thermo Fisher Scientific) and imaged with an FV3000 Confocal Laser Scanning Microscope (Olympus). DAPI or Hoechst 33342 was used for nuclear staining.

1.2 × 10^5^ HEK293-PEX5C11A cells were seeded in 24-well plates on coverslips (Neuvitro #GG1215PLL). After 1day, cells were treated with or without doxycycline (1ug/ml) for 3 days. The cells were rinsed with PBS, fixed in 4% paraformaldehyde for 10 min, and rinsed with PBS again. Cells were permeabilized in 0.5% Triton X-100 in PBS for 10 min. Cells were treated with PBS containing 1% bovine serum albumin (BSA) for 1 hr at room temperature. The cells were incubated with antibody against RPS6 (1:50) diluted in PBS for overnight at 4°C. Next day, cells were washed with 0.05% Triton X-100 in PBS for three times and incubated with Alexa Flour^®^ 488 donkey anti-mouse IgG (1:500) and Hoechst (1:1000) diluted in PBS for 1hr at room temperature in the dark. After that, cells were washed with 0.05% Triton X-100 in PBS for three times. Cells were briefly washed with PBS and mounted on glass slides using mounting medium (Thermo Fisher Scientific, #P36961). Images were visualized by confocal microscope.

### Image analysis and quantification

Confocal images were quantified using Olympus CellSense Dimension software (Olympus, v 1.16) and ImageJ (v1.49). For oenocyte RpS6 signaling quantification, five oenocyte nucleus were randomly selected from each image. Each single nucleus was set as a region of interest (ROI). A background ROI was also selected. The intensity of RpS6 signal was calculated for nucleus intensity and normalized to total intensity (total intensity = nucleus intensity + background). For human RpS6 quantification, we counted the number of cells that that contain RpS6 signal inside the nucleus versus total number of cells in one image.

### Statistical analysis

GraphPad Prism (GraphPad Software, La Jolla, CA, USA, v6.07) or Deseq2 was used for statistical analysis. To compare the mean value of treatment groups versus that of control, student t-test was used. Log2 fold change and FDR values were calculated by Deseq2 using Benjamini-Hochberg method for sequencing data.

---

**Supplementary figure 1:**
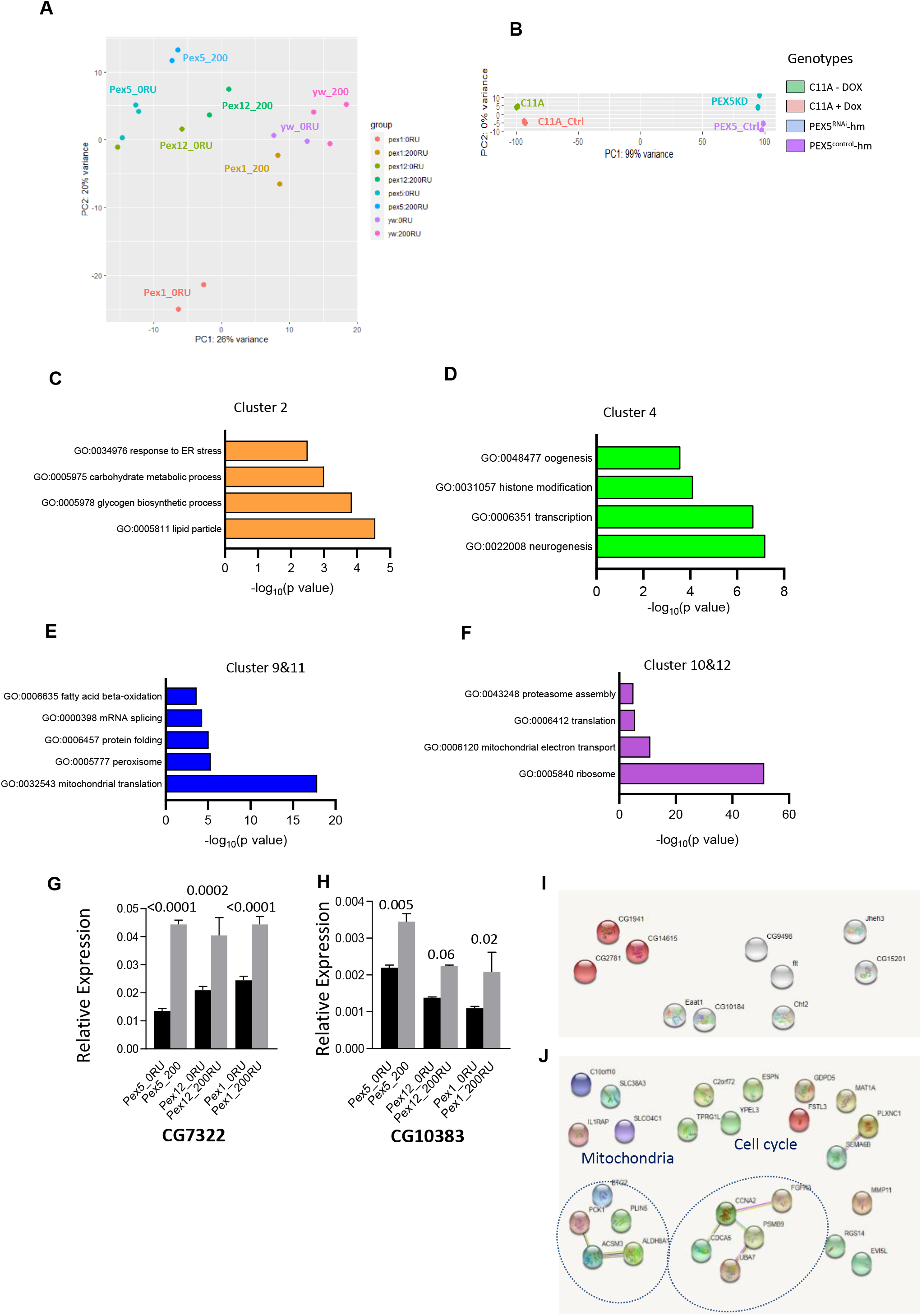
**A** PCA analysis on gene expression from oenocyte samples: Pex1, Pex12, Pex5 and yw. 0RU represents the control group, 200RU represents the RNAi group. **B** PCA analysis on gene expression from human cell culture samples: PEX5^C11A^ (-DOX) versus PEX5^C11A^ (+DOX); control versus PEX5^RNAi^. **C-D** GO analysis for clusters identified in Figure **2A**. **E** Oenocyte genes that are commonly repressed by all Pex knockdowns but not affected by RU feeding. **F** Genes that are commonly induced by PEX5 RNAi and PEX5^C11A^ in human cell cultures.

**Supplementary figure 2:**
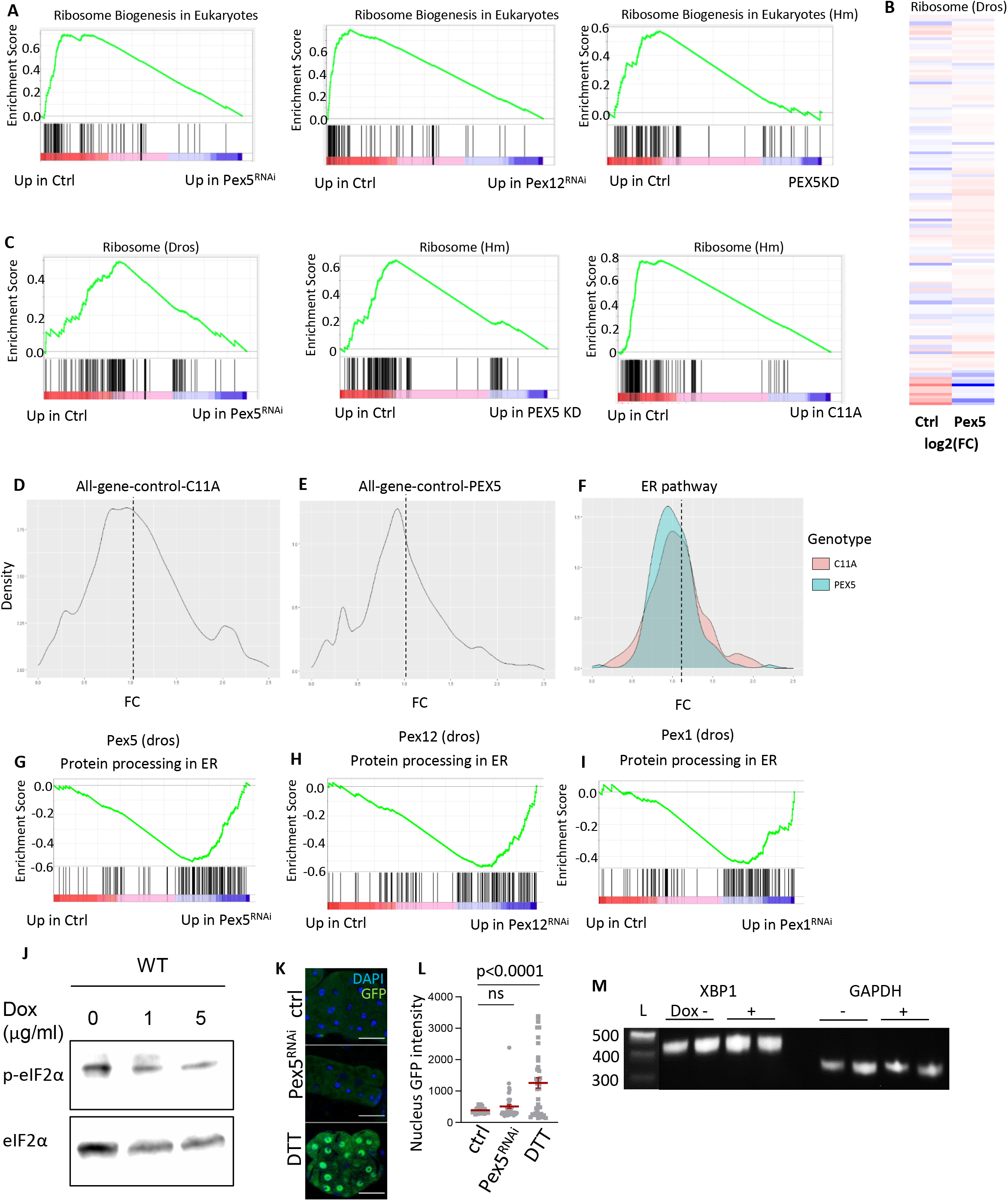
**A** GSEA enrichment profiles on ribosome biogenesis pathway in Pex5 RNAi, Pex12 RNAi of oenocyte samples and PEX5 knockdown in human cell culture. **B** Heatmap analysis on plotting log2 (fold change) value in control group (200 RU/0RU) and Pex5 RNAi samples in oenocytes on ribosome genes. **C** GESA enrichment profiles in ribosome pathway in Pex5 RNAi in oenocyte samples, PEX5 knockdown and PEX5^C11A^ samples in human cells. **D-E** Density plot on fold change of all genes in PEX5^C11A^ group and PEX5 knockdown group. **F** Density plot on fold change of ER pathway genes in PEX5^C11A^ group and PEX5 knockdown group. **G-I** GSEA enrichment profiles on protein processing in endoplasmic reticulum in Pex5 RNAi, Pex12 RNAi and Pex1 RNAi samples in oenocytes. **J** Western blot measuring P-eIF2α and eIF2α level in control cell line treated with increasing concentration of Doxycycline (0 ug/ml, 1 ug/ml, and 5 ug/ml). N = 2. **K** Immunostaining of oenocytes expressing xbp1-EGFP reporter with anti-GFP. EGFP is out of frame without Ire-1-mediated splicing but comes in frame after splicing. Xbp1-EGFP marker is activated by Ire-1 in response to DTT treatment (bottom panel). The middle panels show immunostaining of oenocytes with control or *Pex5* knockdown. **L** Quantification on nucleus GFP intensity in **K**. **M** PEX5^C11A^ cells were treated with Doxycycline (5 ug/ml) or without, and cell lysates were obtained. RNA extracted was subjected to RT-PCR with *XBP1* primers. The lengths of DNA size markers are shown on the left.

