## Supplemental Table 1 for "Peroxisome import stress impairs ribosome biogenesis and induces integrative stress response through eIF2α phosphorylation"

*Supplementary Table 1: Primer list*

| Oligonucleotides | Sequence 5'-3' | Species |
| --- | --- | --- |
| ITS1-F | TTATTGAAGGAATTGATATATGCC | Drosophila |
| ITS1-R | ATGAGCCGAGTGATCCAC | Drosophila |
| ETS-F | GCTCCGCGGATAATAGGAAT | Drosophila |
| ETS-R | ATATTTGCCTGCCACCAAAA | Drosophila |
| ETS-F-unprocessed | cgagtgctatataaaaatggccg | Drosophila |
| ETS-R-unprocessed | gcatataactactggcaggatc | Drosophila |
| 18S-f | TGGTCTTGTACCGACGACAG | Drosophila |
| 18S-r | GCTGCCTTCCTTAGATGTGG | Drosophila |
| ETS-F-unprocessed | ctcgccgcgctctacctt | Homo Sapiens |
| ETS-R-unprocessed | gcgcccgtcggcatgtattagctc | Homo Sapiens |
| 18S-f | CTTTCGATGGTAGTCGCCGT | Homo Sapiens |
| 18S-r | CCTTGGATGTGGTAGCCGTT | Homo Sapiens |
